# Interplay between extreme drift and selection intensities favors the fixation of beneficial mutations in selfing maize populations

**DOI:** 10.1101/2020.12.22.423930

**Authors:** Arnaud Desbiez-Piat, Arnaud Le Rouzic, Maud I. Tenaillon, Christine Dillmann

## Abstract

Population and quantitative genetic models provide useful approximations to predict long-term selection responses sustaining phenotypic shifts, and underlying multilocus adaptive dynamics. Valid across a broad range of parameters, their use for understanding the adaptive dynamics of small selfing populations undergoing strong selection intensity (thereafter High Drift-High selection regime, HDHS) remains to be explored. Saclay Divergent Selection Experiments (DSEs) on maize flowering time provide an interesting example of populations evolving under HDHS, with significant selection responses over 20 generations in two directions. We combined experimental data from Saclay DSEs, forward individual-based simulations, and theoretical predictions to dissect the evolutionary mechanisms at play in the observed selection responses. We asked two main questions: How do mutations arise, spread, and reach fixation in populations evolving under HDHS ? How does the interplay between drift and selection influence observed phenotypic shifts ? We showed that the long-lasting response to selection in small populations is due to the rapid fixation of mutations occurring during the generations of selection. Among fixed mutations, we also found a clear signal of enrichment for beneficial mutations revealing a limited cost of selection. Both environmental stochasticity and variation in selection coefficients likely contributed to exacerbate mutational effects, thereby facilitating selection grasp and fixation of small-effect mutations. Together our results highlight that despite a small number of polymorphic loci expected under HDHS, adaptive variation is continuously fueled by a vast mutational target. We discuss our results in the context of breeding and long-term survival of small selfing populations.

---

Understanding the evolutionary processes sustaining phenotypic shifts is at the core of quantitative genetic models. Empirical description of such shifts takes its roots in the breeding literature where truncation selection generates significant and sustainable responses (Hill and Caballero 1992; Walsh and Lynch 2018). Truncation selection is known to be the most effective form of directional selection (Crow and Kimura 1979). Under truncation selection, limits to the evolution of phenotypes are rarely reached as heritable variation persists through time (Odhiambo and Compton 1987; Moose *et al.* 2004; Weber and Diggins 1990; Caballero *et al.* 1991; Mackay 2010; Lillie *et al.* 2019). Such observations fit well with the breeder equation and its derivatives (Lush 1943; Lande 1979; Lande and Arnold 1983) which accurately predict selection response after one generation. With the additional hypothesis of constant genetic variance provided by the Fisher’s infinitesimal model (Fisher 1930), theoretical models predict a continuous and linear response with no finite limits. However, the rate of response is expected to decline with selection-induced linkage disequilibrium (Bulmer 1971; Hospital and Chevalet 1996). Furthermore under finite population size, selection response is predicted to reach an asymptotic finite limit (Robertson 1960) as exemplified in mice (Roberts 1967; Falconer 1971). Results from other species are more equivocal (e.g. drosophila (Weber 1990; Weber and Diggins 1990; Weber 1996), or maize (Odhiambo and Compton 1987; Moose *et al.* 2004; Dudley and Lambert 2010; De Leon and Coors 2002; Lamkey 1992). Incorporation of *de novo* mutations indeed predicts a slower rate of response instead of a hard limit (Hill 1982b,a; Weber and Diggins 1990; Wei *et al.* 1996; Walsh and Lynch 2018). A sub-optimal average selection response is expected in two situations: when population size, *N* is below 10^4^ reducing the genetic variance 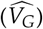 at mutation-drift equilibrium(Hill 1982b; Houle 1989); and when 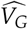 is reduced due to strong selection (Houle 1989). Overall, quantitative genetic models that include selection, drift and mutation (Houle 1989) are well-suited for predicting observed selection responses in a broad range of parameters (Hill and Rasbash 1986) — providing appropriate corrections, *e.g.* deviations from low drift and low selection intensity (Walsh and Lynch 2018). Most of these models, however, make the assumptions of random mating and of a probability of fixation of new mutations — determined by the product of population size by their selection coefficient, *N_s_* — to be either ≪ 1 or ≫ 1. Mathematical models for the intermediate regime *N_s_* ≈ 1 and non-random mating still remain unsatisfactory. Furthermore, the description of mechanisms of long-term selection response — and whether it can be understood and predicted by existing equations — has yet to be explored for polygenic traits evolving in small selfing populations under high selection intensity, a regime subsequently called HDHS (High-Drift High-Selection).

Both the Distribution of mutational Fitness Effects (DFE) and the mutation rate are central to long-term predictions of selection responses. Selection makes the DFE of fixed mutations different from that of incoming *de novo* mutations (Kassen and Bataillon 2006). In large populations, a high proportion of incoming *de novo* beneficial mutations are predicted to reach fixation, together with vanishing small effect deleterious mutations (Crow and Kimura 1971; Kimura 1983). In small populations and/or at small selection intensity, frequent loss of beneficial mutations due to drift together with the fixation of moderately strong deleterious mutations is expected. Hence Kimura’s equation that links the fixation probability (*P_fix_*) of a mutation to its frequency (*p*), the population size (*N*) and selective coefficient (*s*) — *P_fix_* (*s*, *p*, *N*) = (1 − *e*^−4*spN*^)/(1 − *e*^−4*sN*^) — applies to a vast range of parameters including *s* values as high as 0.1 and *N* as small as 10 individuals (Carr and Nassar 1970). An additional layer of complexity to DFE prediction comes from the mating system. Adaptation of very large asexual populations (such as microbes) is indeed affected by competition between alternative beneficial mutations occurring in different genetic background, a process referred to as clonal interference (Gerrish and Lenski 1998). Here the absence of recombination favors enrichment of the DFE in large beneficial mutational effects (Gerrish and Lenski 1998). However, if selection overpowers drift, *i.e. Ns* ⪆ 1, or if the rate of beneficial mutation (*μ_B_*) is small enough, the expected time lag between two successive mutations is sufficiently large for the first beneficial mutation to fix without interference of the second. While such behavior is expected when *Nμ_B_* ≪ 1/*ln*(*N_s_*), for *Nμ_B_* ⪆ 1/*ln*(*N_s_*) beneficial mutations evolve under clonal interference (Desai and Fisher 2007). Altogether these results highlight how the interplay of key parameters - *N*, *s*, *μ*, effective recombination — determine the DFE and in turn, the long-term selection response.

Genomic footprints of selection have considerably enriched our vision of allele trajectories sustaining selection responses. On the one hand, one can observe genomic footprints such as hard and/or soft selective sweeps. A hard sweep is characterized by a strong decrease in genomic diversity at the selected locus and its surrounding region through genetic hitchhiking (Hermisson and Pennings 2017); while a soft sweep is associated with a weak genomic signature either because recombination on standing variation occurs so that a given advantageous mutation is associated with multiple haplotypes, or because recurrent *de novo* mutations are associated with multiple haplotypes. Together these footprints indicate that adaptation proceeds through a succession of sweeps at loci encoding the trait. On the other hand, absence of selection footprints is expected under the so-called polygenic selection model (Berg and Coop 2014; Wellenreuther and Hansson 2016; Walsh and Lynch 2018), that rather posits a collective response at many loci translating into simultaneous subtle shifts in allele frequencies, in compliance with the infinitesimal model. Whether adaptation proceeds through hard/soft sweeps or polygenic model primarily depends on the population-scaled mutation rate (*θ*) as well as the number of redundant loci that offer alternative ways for adaptation (*L*) — the mutational target. Adaptation proceeds through hard sweeps for small *θ × L* (≤0.1) while polygenic adaptation requires large *θ × L* (≥ 100) with partial/soft sweeps in between (Messer and Petrov 2013; Höllinger *et al.* 2019). Extension of the hitchhiking model to a locus affecting a quantitative trait with an infinitesimal genetic background predicts that, under the hypothesis of a Gaussian fitness function, the fixation of a favorable mutation critically depends on the initial mutation frequency and the distance to the optimum (Chevin and Hospital 2008). Interestingly, while demographic parameters— population size, bottleneck strength — play a relatively small role in the speed of adaptation compared to standing and mutational variance, they change its qualitative outcome. Population bottlenecks diminish the number of segregating beneficial alleles, favoring hard sweeps from *de novo* mutations over soft sweeps from standing variation (Stetter *et al.* 2018).

By exploring short-term temporal dynamics of adaptation, experimental evolution has provided further hints into allele frequency changes, and into the extent of polymorphism and competition among beneficial mutations under various drift/selection/recombination regimes. Temporal dynamics are obtained either through pedigree information or time series samples. This last approach, widely used in microorganisms has revealed complex patterns of mutation spreading during the course of adaptation. These include clonal interference, the reduction of the relative advantage of a beneficial mutation in fit *versus* less fit genotypes (diminishing-return epistasis), and evidence for the same favorable mutation being selected in multiple independent evolved clones (genetic parallelism) (Good *et al.* 2017; Spor *et al.* 2014; Neher 2013; Good *et al.* 2012; Desai and Fisher 2007; Gerrish and Lenski 1998). However, in asexually reproducing microbes, adaptation proceeds through *de novo* mutations, which may reveal specific patterns not found in sexually-reproducing eukaryotes. In yeast, for instance, most adaptive changes correspond to the fixation of initial standing variation (Burke *et al.* 2014; Burke 2012). Patterns of allele frequency changes depend crucially on both *N_e_* and the frequency of sex, that are themselves intimately linked (see Hartfield *et al.* (2017)). Considering a single locus, fixation time decreases correlatively with the level of self-fertilization (Haldane 1927). At the same time, multilocus simulations have shown that selfing reduces effective population size through background selection and in turn, beneficial mutations are less likely to fix (Kamran-Disfani and Agrawal 2014; Roze 2016). In addition, as selection interference reduces the efficiency of selection in low-recombining regions, high selfing rates also increase the fixation of deleterious mutations through genetic hitchhiking (Hartfield and Glémin 2014). These insights are together in line with the low selection approximation that posits that reduction in effective recombination decreases selection efficiency.

In the current paper, we aimed at investigating the dynamics of the response to selection in small selfing populations evolving under high selection intensity. Situated at the parameters boundaries of current models, this regime is of particular interest to understand the limits of adaptation and long-term survival of small selfing populations undergoing strong selection. We relied here on two Divergent Selection Experiment (DSEs) conducted for 18 generations on Saclay’s plateau (Saclay DSEs), south of Paris (France). These Saclay DSEs are ideal settings to address those issues: selection-by-truncation has been applied in a higher organism (maize), on a highly polygenic and integrated trait (flowering time, (Buckler *et al.* 2009; Tenaillon *et al.* 2018)) that directly affects fitness. Previous results indicate continuous phenotypic responses sustained by a constant mutational input (Durand *et al.* 2010, 2012, 2015) — values of mutational heritability ranged from 0.013 to 0.025. We asked two main questions: How do mutations arise, spread, and reach fixation in populations where both drift and selection are extremely intense ? How does the interplay between drift and selection influence the response to selection ? To answer those questions, we confronted the observed phenotypic responses in Saclay DSEs to forward individual-based simulations that explicitly modeled the same selection and demographic scheme, and used theoretical predictions to measure deviations from expectations.

## Materials and Methods

### Saclay Divergent selection experiments

We have conducted two independent divergent selection experiments (Saclay DSEs) for flowering time from two commercial maize inbred lines, F252 and MBS847 (thereafter MBS). These experiments were held in the field at Université Paris-Saclay (Gif-sur-Yvette, France). The selection procedure is detailed in Fig. S1 and Durand *et al.* (2010). Briefly, within each Saclay DSE, the ten earliest/ten latest flowering individuals were selfed at each generation to produce 100 offspring used for the next generation of selection within the Early/Late populations, so that 1000 plants were evaluated in each population. Following Durand *et al.* (2015), we designated as *progenitor*, a selected plant represented by its progenies produced by selfing and evaluated in the experimental design at the next generation. Seeds from progenitors from all generations were stored in cold chambers.

Within each population, we evaluated offspring of a given progenitor in four rows of 25 plants randomly distributed in a four-block design. Each block contained 10 rows representing the 10 progenitors. We applied a multi-stage pedigree selection. First the three earliest (latest) flowering plants within each row were selfed and their flowering time was recorded. This corresponded to 12 plants per progenitor, *i.e.* 120 plants per population. The second stage consisted in choosing 10 plants among the 120 on an index based on three criteria : flowering time, total kernel weight and pedigree. When two plants had the same flowering date, we chose the one with the highest kernel weight. In addition, we maintained two independent *families* within each population, *i.e.* two sub-pedigrees derived from two different progenitors in the ancestral *G*_0_ population, and we never selected more than three plants from the same *G_n − 1_* progenitor. Practically, each family was composed of three to seven progenitors at each generation. Altogether, we selected in each population 10 plants out of 1000 which corresponded to a selection intensity of 1%.

We traced back the F252 and MBS pedigrees from generation 20 (*G*_20_) to the start of the divergent selection experiments, *G*_0_. The initial MBS pedigrees encompassed four families: ME1 and ME2 for the MBS Early (ME) population, and ML1 and ML2 for the MBS Late (ML) population (Fig. S2). F252 Early (FE) population was composed of FE1 and FE2 families (Fig. S2). F252 Late populations genealogies were more complex: FVL families (F252 Very Late in Durand *et al.* (2015)) ended at generation 14 with the fixation of a strong effect allele at the *eIF-4A* gene (*Durand et al.* 2015). To maintain two families in F252 Late population, two families FL2.1 and FL2.2 were further derived from the initial FL2. These two families pedigrees are rooted in FL2 from a single *G*_3_ progenitor (Fig. S2).

### Phenotypic data collection and empirical selection responses

The same approach as Durand *et al.* (2015) was applied. Briefly, progenitor flowering dates, measured here as the number of days to flowering after sowing equivalent to 20°C days of development (Parent *et al.* 2010), were recorded as the 12 earliest or latest plants in their progeny at each generation of the Saclay DSEs. We used these records to investigate the response to selection treating each family independently. After correction of the phenotypic values *Z_ijklmn_* for block effects, and year effects according to equation (1) of Durand *et al.* (2015) (so that 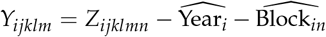, *in* corresponding to the block effect *n* in selection year *i*), the linear component *b_jk_* of the within-family response to selection was estimated using the following linear model:

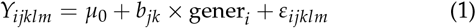

where *i* stands for the year and corresponding generation of selection (so that gener_*i*_ takes values between 0 to 20) *j* for the population (Late or Early), *k* for the family within population (*e.g* ME1 or ME2), *l* for progenitor within family, and *m* for the plant measurements within progenitor (so that *ε_ijklm_* corresponds to the residual variance due to differences between progenitor of the same generation, and family and plant effect). Finally, *μ*_0_ is the intercept corresponding to the average flowering time at generation *G*_0_,

Family means and standard errors were also computed at each generation to represent families selection responses presented Fig. 1 (a). All the values were centered around 100 for comparison purposes with the simulated responses.

**Figure 1.**
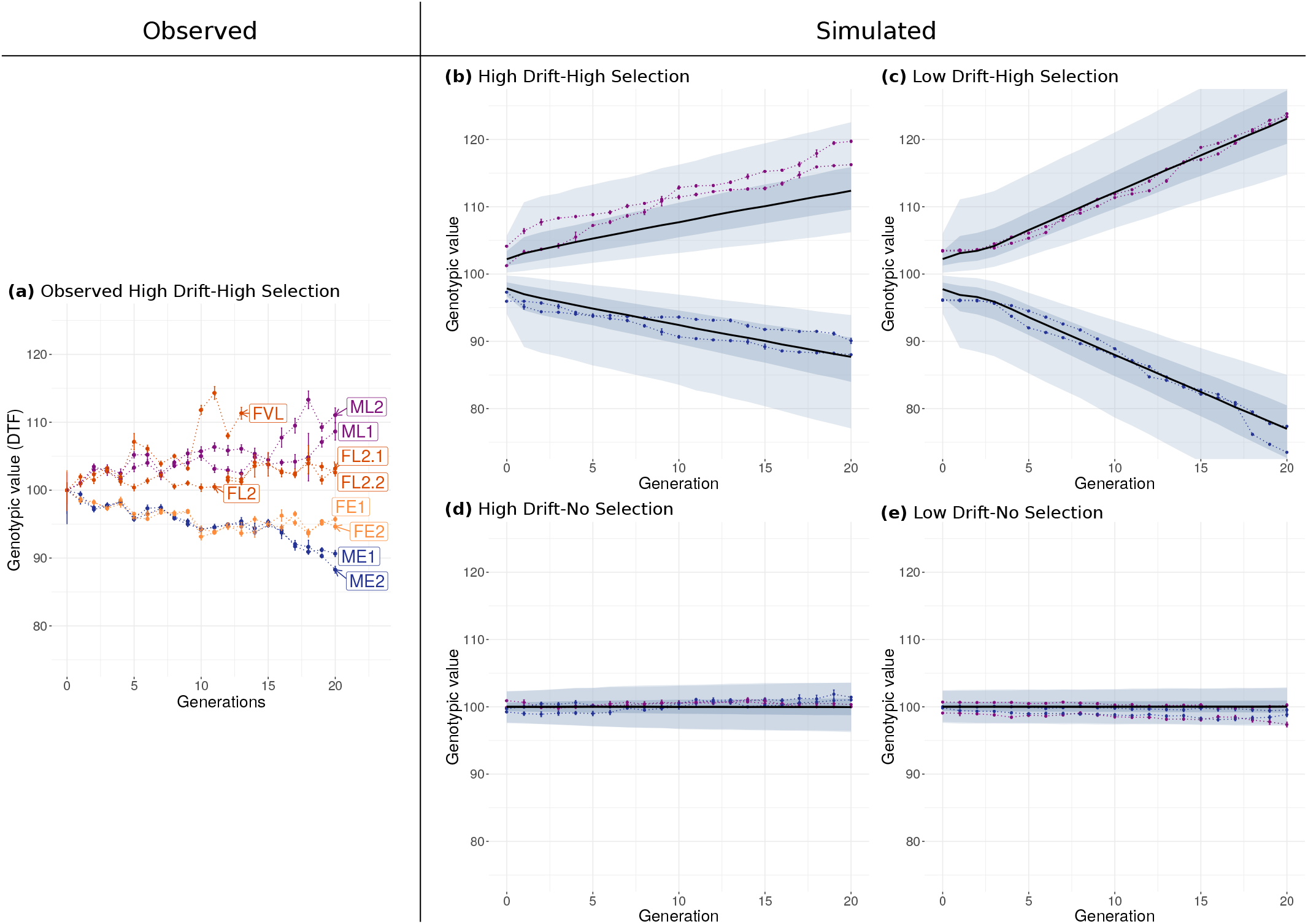
Observed and simulated selection response. Selection response is visualized by the evolution of the mean genotypic values of the selected progenitors per family (expressed in Days To Flowering, DTF) across generations in observed (a) and simulated (b-e) data. Observed genotypic values correspond to mean phenotypic values corrected for environmental effects. In (a), red/orange corresponds to late/early flowering F252 families, while violet/blue corresponds to late/early flowering MBS families. All families were centered around 100, and Vertical bars correspond to ±1 genotypic standard error around the mean. We simulated four regimes with the parameters calibrated from the MBS observed response: High Drift-High Selection intensity (b), Low Drift-High Selection (c), High Drift-No Selection (d), Low Drift-No Selection (e). Violet/blue color identifies late/early population. In each population, the black line represents the evolution of the median value over 2000 simulations of the family genotypic mean. The shaded area corresponds to the 5^*th*^-95^*th*^ percentiles (light blue) and to the 25^*th*^-75^*th*^ percentiles (dark blue). In addition, two randomly chosen simulations are shown with dotted lines

### Model framework

We used forward individual-based simulations that explicitly modeled the same selection — proportion of selected individuals=1% of the most extreme) — and demographic scheme — variations in population size — as Saclay DSEs. This regime is referred to High-Drift High Selection intensity (HDHS). **Initial *G*_0_ simulation**: We obtained our initial population by mimicking a classical selection scheme used to produce fixed maize inbred lines in industry. To do so, we started from an heterozygous individual that was selfed for eight generations in a single-seed descent design. An additional generation of selfing produced 60 offspring that were reproduced in panmixia for two generations to constitute the 60 individuals of the *G*_0_ initial population. Therefore, we started our simulations with a small initial residual heterozygosity (≤ 0.5%). ***G*_1_ simulation**: Considering one Saclay DSE, we selected from the initial population (60 individuals), the two earliest and the two latest flowering parents on the basis of their average phenotypic value. Each of these individuals constituted the ancestor of each of the four families. They were selfed to produce 100 offspring. **Subsequent generations *n***: From there, we simulated the exact same selection scheme that included a two-steps procedure (Fig. S1). First, we selected the 12 earliest (within each early family) and the 12 latest individuals (within each late family) from the 100 offspring of each progenitor. We next selected the five earliest (within each early family) and five latest (within each late family). In other words, at each step we retained 5 out of 500 (5/500) individuals within each of the four families. Note that we imposed that the five selected individuals did not share the same parent.

### Simulated genetic and phenotypic values

Maize flowering time is a highly polygenic trait (*Buckler et al.* 2009; Tenaillon *et al.* 2018). Over 1000 genes have been shown to be involved in its control in a diverse set of landraces (Romero Navarro et al. 2017). We therefore set the number of loci to *L* = 1000. As in maize, the genome of one individual was composed of 10 chromosomes. In each simulation: (i) we randomly assigned each locus to a chromosome so that genome composition varied from one simulation to another; (ii) the position of each locus within each chromosome was uniformly drawn between 0 and 1.5, 1.5 Morgan being the total genetic length of each chromosome; (iii) the crossing-over positions along chromosomes were drawn in an exponential law of parameter 1, which corresponded to an effective crossing-over every Morgan. The initial population (*G*_0_) consisted of 60 individuals polymorphic for a small fraction of loci (residual heterozygosity). Let 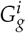 be the genotype of the individual *i* of the generation *g*. Let 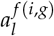 the allelic value at the locus *l* of the paternal chromosome *f* of the individual *i* at the generation *g* and 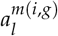 the allelic value at locus *l* of maternal chromosome *m* of individual *i* at generation *g*. This allows us to model the genotype of an individual as :

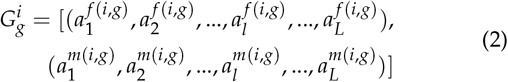

We expect the distribution of allele effects to follow a leptokurtic Gamma distribution (*e.g.* Kimura (1979); Hill (1982a); Keightley (1994); Shaw *et al.* (2002); Piganeau and Eyre-Walker (2003)). We made the simplifying assumption that the unknown shape parameter of the Gamma distribution *α* was equal to 1, corresponding to an exponential distribution. Overall, the initial allelic values were drawn in an reflected exponential distribution, that is to say:

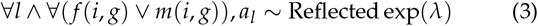

Hence the probability density:

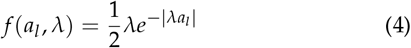

which implied that:

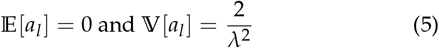

Starting from a hybrid heterozygote at all *L* loci, we computed the expectation of the genic variance 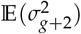 after *g* generations of selfing and two generations of bulk. Selfing reduces the genic variance by 1/2 each generation. In the absence of linkage disequilibrium, panmixia does not change allelic frequencies:

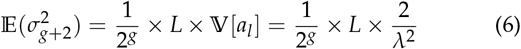

Therefore, to match the field estimate 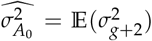, one could let

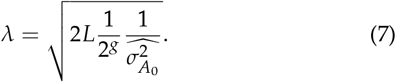

However, drift, linkage disequilibrium and mutation can lead to deviations from the expected value of the initial genetic variance. We therefore recalibrated all the allelic values at generation 0 to match the initial 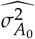 additive variance. To do so, we multiplied all the allelic values by a corrective factor 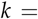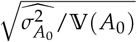, where 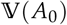 was the additive variance of our population *G*_0_, calculated in multiallelic as 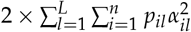 with *n* the number of alleles at locus *l*, *p_il_* the frequency of the allele *i* at locus l. *α_il_*, its additive effect, is defined as in Lynch *et al.* (1998) (Chapter 4), so that after dropping subscript 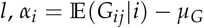, with *μ_G_* the population genotypic mean, 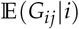 the conditional expectation of the genotypic value of genotype *G_ij_* knowing *i*. So at *G_0_*, 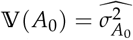.

Mutations occurred at each reproduction event. We drew the number of mutations per haplotype in a Poisson distribution of mean *L × μ* where *μ* was the mutation rate per locus. Following Kimura (1979); Hill (1982a); Keightley (1994); *Shaw et al.* (2002); Piganeau and Eyre-Walker (2003), we drew the value of a mutation at a locus in a reflected exponential distribution of parameter 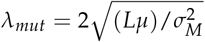. We computed phenotypic values as the sum of all allelic values *a_il_* (*L* × 2) plus an environmental effect randomly drawn in a normal distribution of mean 0 and variance 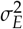.

### Selection and drift regimes

As control, we considered a model without selection (the No Selection regime, NS) where neutral evolution occurred in a population with the same census size and the same number of progenitors as in the selection model. At each generation, we randomly drew 1% of the individuals to form the next generation, instead of choosing them from their phenotypic values.

In addition, we considered an alternative drift regime, where we increased the census population size by a factor 10, all other parameters remaining unchanged. This regime is referred to as the Low Drift regime (LD) where 50/5000 instead of 5/500 individuals were selected. Both Low Drift-High Selection (LDHS) and Low Drift-No selection (LDNS) were considered.

We performed 2000 independent simulations for each of the four families in each of the four regimes. All downstream analyses were carried out over all simulations, except when specified.

### Parameter calibration

Our model encompassed three key parameters: the initial additive variance 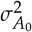, the mutational variance, 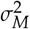, and the environmental variance 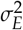. We chose to sample parameter values in an inverse-gamma distribution with parameters (shape and scale) chosen such as (i) the expected means of the two inverse-gamma for 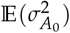 and 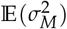 were roughly equal to the estimate provided by (Durand *et al.* 2010) for MBS-DSE, (ii) 95% of the values of the two inverse-gamma fell within the range of observed values for the DSEs (Durand *et al.* 2010), (iii) mean and variance for 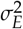 corresponded to the values measured for maize experiments in Saclay’s Plateau. Parameters values are summarized Tab. 1.

**Table 1.** Initial variances parameters.

| Parameter | Expectation | Variance | shape | rate |
| --- | --- | --- | --- | --- |
| $\sigma_{A_0}^2$ | 2.25 | 3.28 | 3.540 | 5.715 |
| $\sigma_M^2$ | $3.38 \times 10^{-2}$ | $7.27 \times 10^{-5}$ | 17.67 | 0.5626125 |
| $\sigma_E^2$ | 2.25 | 0.32 | 17.67 | 37.50075 |

Genomic parameters were taken from the literature. In maize, Clark *et al.* (2005) estimated the nucleotidic substitution rate to 30 × 10^−9^. We relied on the maize reference genome V4 (Jiao *et al.* 2017) to estimate an average mRNA length of 6000 (median=5197, mean=7314). Based on both estimates, we therefore considered a mutation rate per locus of: *μ* = 6000 × 30 × 10^−9^ = 1.8 × 10^−4^.

### Expected response, effective population size and time to the most recent common ancestor

We computed the expected cumulative response after *t* generations for haploid population as (Hill 1982b; Wei *et al.* 1996; Weber and Diggins 1990; Walsh and Lynch 2018):

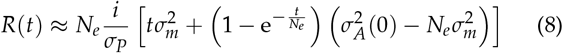

The effective population was the only parameter not explicitly defined in our simulations and is of crucial importance in the response to selection. We estimated *N_e_* following two approaches. First using the Time to the Most Recent Common Ancestor (TMRCA) from the standard coalescence theory for a haploid sample of size *k* at generation *g* (Walsh and Lynch 2018):

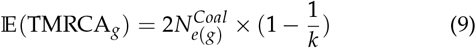

Second, from the variance in offspring number (Crow and Kimura 1971; Durand *et al.* 2010), where *N_e_* can be computed as

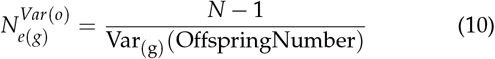

In the simulations, 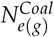 and 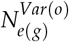 were computed at generation *G*_20_. We also computed the harmonic means between generations *G*_1_ and *G*_20_ and computed the whole distribution (in 2*N_e_* generations) of the Kingman coalescent TMRCA as (Tavaré 1984):

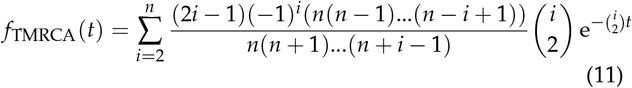

### Fitness function and Kimura’s expected fixed mutational DFE

Using diffusion equations, Kimura (Kimura 1962) predicts the fixation probability of a mutation of selective value *s*(*a*) — with *a* its allelic value — and initial frequency *p*:

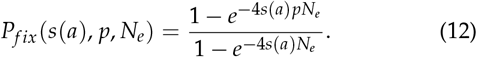

When occurring, a new mutation arises during meiosis in one plant among the 500 of a family observed at a given generation. Hence, its effect on the phenotypic variance is negligible. Therefore, the plant carrying this newly arisen mutation was selected essentially independently of the new mutation. Consequently, the 5 individuals selected in one family comprised one heterozygote (Aa) bearing the mutation, and 4 homozygotes (aa). Each selected individual produced 100 progenies, so that the fitness effect of the mutation was evaluated at the next generation in a population of 500 plants where the frequency of the mutant allele was *p* = 1/10 (Table 2).

**Table 2.** Fitness model.

| Genotype | AA | Aa | aa |
| --- | --- | --- | --- |
| Genotypic frequency | 1/20 | 2/20 | 17/20 |
| Fitness value | $w_{AA}$ | $w_{Aa}$ | $w_{aa}$ |
| Mutational effect | $a_{AA} = 2a$ | $a_{Aa} = a$ | $a_{aa} = 0$ |

In this population, the distribution of flowering time resulted from a mixture of Gaussian distributions.

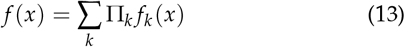

where *f_k_* (*x*) is the flowering time distribution for plants with genotype *k ∈ AA*, *Aa*, *aa*. As we selected 1% of the latest/earliest flowering plants, all selected plants did flower after the date *z*, computed as the 1% quantile of the mixture distribution. The selection effect *s*(*a*) depended on the effect *a* of the mutation on flowering time (Table 2). Indeed, the relative weight of homozygous mutants *AA* among selected individuals was computed as:

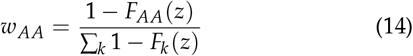

Which leads to:

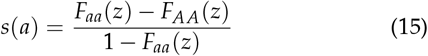

The fixation probability *P_fix_* (*s*(*a*), *p*, *N_e_*) was computed as in (Eq. 12) using *s*(*a*) (Eq. 15), *p* = 1/10, and 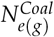 for *N_e_*. The mutational effect *a* was drawn in a reflected exponential distribution of parameter *λ_mut_* and density function *g_λmut_* (*a*). Hence, the density of fixed mutations *h*(*a*) was computed as:

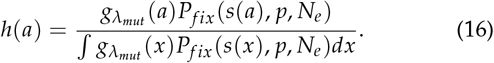

Moreover, we recorded the simulated values *a_sim_* of each fixed mutation and computed the realized distribution *h_obs_* (*a*) using kernel estimate methods.

## Results

In order to examine the evolution and fate of small selfing populations submitted to strong selection intensity, we investigated the dynamics of the response to selection under a High Drift-High Selection intensity (HDHS) regime imposed on two divergent artificial selection experiments for flowering time in maize (Saclay DSEs). We compared experimental data to results of a simulation model specifically devised to mimic our experiments; and further computed when possible expectations from population and quantitative genetics theory.

### Empirical response after 20 generations of selection

In line with previous observations for the first 16 generations, we observed significant responses (Fig. 1 a, Tab. 3a, 3b) to selection after 20 generations in all families. Marked differences among families nevertheless characterized these responses. This is well exemplified in the Late F252 families where one family (FVL) responded very strongly with a mean shift of 11.32 Days to Flowering (DTF) after 13 generations, corresponding to a linear regression coefficient of 0.86 DTF/generation (Tab. 3a). This family fixed a deleterious allele at *G*_13_ and could not be maintained further (Durand *et al.* 2012). We examined two derived families from *G*_3_ (Fig. S2), the FL2.1 and FL2.2. These families were shifted by 3.19 DTF and 2.60 DTF from the *G*_0_ FL2 mean value for FL2.1 and FL2.2, respectively. These corresponded to a linear regression coefficient of 0.11 DTF/generation for FL2.1 and 0.12 DTF/generation for FL2.2 (Tab. 3a). The selection response were more consistent for the two Early F252 families, with a shift after 20 generations of −4.27 DTF for FE1, and a shift of −5.34 DTF for FE2 (Tab. 3a). Considering MBS genetic background, the late/early MBS families were shifted by 8.64 DTF for ML1, and 11.05 DTF for ML2 (respectively −9.34 DTF for ME1 and −11.72 DTF for ME2), with linear regression coefficient of 0.24 DTF/generation for ML1, and 0.46 DTF/generation for ML2 (respectively −0.41 DTF/generation for ME1 and - 0.42 DTF/generation for ME2) DTF (Tab. 3b).

**Table 3.**
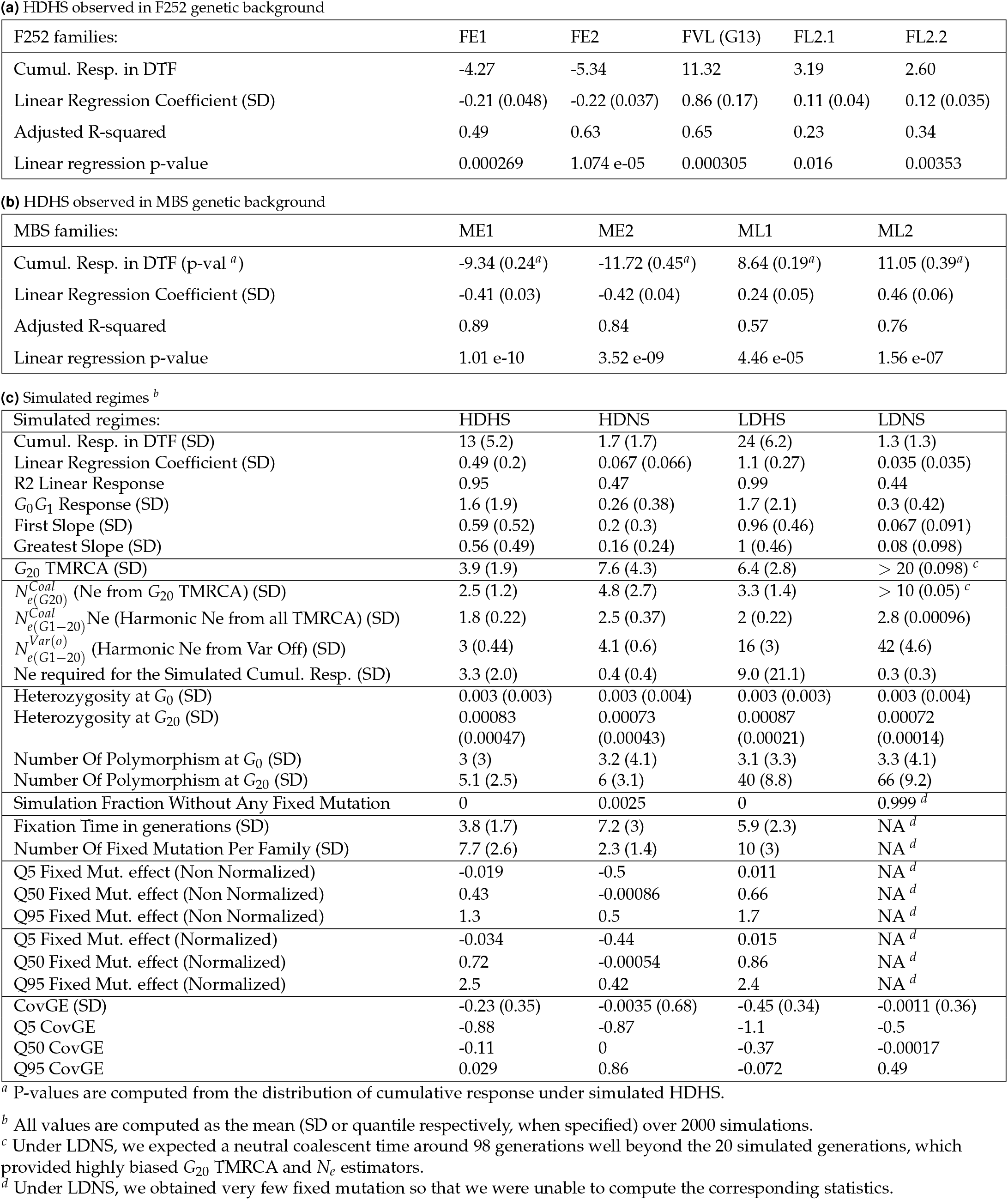
Descriptive statistics of the selection response dynamics in observed F252 genetic background (a), observed MBS genetic background (b) and the 4 simulated regimes (c).

### Simulation model validation

To parameterize our simulation model, we used priors: variance components were described by inverse-gamma distributions whose parameters were chosen following previously reported values for Saclay’s DSEs Durand *et al.* (2010), mutation rate was taken from the maize literature (Clark *et al.* 2005), and the number of loci was set to *L* = 1000 according to the large mutational target described for maize flowering time (Romero Navarro *et al.* 2017). In order to validate the parametrization of our model, we compared the observed MBS responses in all families to the simulated selection responses under HDHS regime. Because of the symmetry in the model construction and for simplicity, simulated results are described for late populations only. We recovered a simulated response with a mean genetic gain of 0.49 DTF/generation (Fig. 1, Tab. 3c). Starting from a mean genotypic value of 100 DTF, the mean genotypic value was shifted by 13.0 DTF (SD: 5.2) after 20 generations. Our simulated response therefore closely matched the observed response (p-value not significant, Tab. 3b) indicating an accurate parametrization of our simulation model (Fig. 1), that captured the average selection response per generation of MBS. Note however, that inter-generational fluctuations were higher in the observations than in the simulations (Fig. 1).

We used simulations both to validate our model and to explore two drift intensities, High and Low. We used corresponding negative controls with No Selection (NS) which lead to four regimes: High Drift-High Selection intensity (HDHS, the default regime), High Drift-No Selection (HDNS), Low Drift-High Selection intensity (LDHS) and Low Drift-No Selection (LDNS).We formally tested the significance of our simulated response by comparing the linear response under HDHS to that obtained under HDNS. We were able to reject the null hypothesis of no selection response in 96.4% of the simulations under HDHS (P-value<0.05).

To investigate the impact of a ten-fold increase of the census population size on selection response, we contrasted HDHS to LDHS. Just like for HDHS, we obtained a significant response under LDHS with a mean genetic gain of 1.10 DTF/generation (Fig. 1, Tab. 3c). This gain was greater than the +0.035 DTF/generation (SD: 0.035) obtained for the LDNS control model, and we were able to reject the null hypothesis of no selection response in 100% of the simulations. The gain under LDHS corresponded to a shift of +24 DTF (SD: 6.2), which was substantially higher than that observed under HDHS. Hence multiplying the census population size of HDHS by 10 (LDHS) resulted in roughly doubling the selection response.

In sum, we validated the accuracy of our model by showing that the simulated response closely matched the observed response. We further demonstrated that selection triggered the response in all populations under both Low and High Drift. Finally, we confirmed our expectation that the selection response was higher in a Low Drift than in a High Drift regime.

### Effective population size

We estimated coalescent effective population sizes *N_e_* from the standard coalescence theory (Eq: (9)) using a Wright-Fisher population of size 5 (HD) and 50 (LD) individuals. With 5 individuals, we expected a theoretical coalescence time around 8 generations, and with 50 individuals, around 98 generations (i.e. more than the number of simulated generations). Focusing on the last generation, our simulations provided estimates of the mean *G*_20_ TMRCA of 7.6 generations under neutrality (NS) for HD, closely matching the theoretical expectation of 8 (Tab. 3c). Considering the LDNS simulations, theoretical expectations (98) largely exceeded the number of generations (20). In contrary, we found a mean *G*_20_ TMRCA of 3.9 under HDHS, and 6.4 under LDHS. Fig. 2 shows the distribution of the TMRCA estimated at *G*_20_ in the three regimes. Under HDNS, the distribution fits the expectation from Eq: (11). As compared to the neutral case, Fig. 2 also shows that both the high drift (HDHS) and low drift (LDHS) selection regimes display reduced TMRCA.

**Figure 2.**
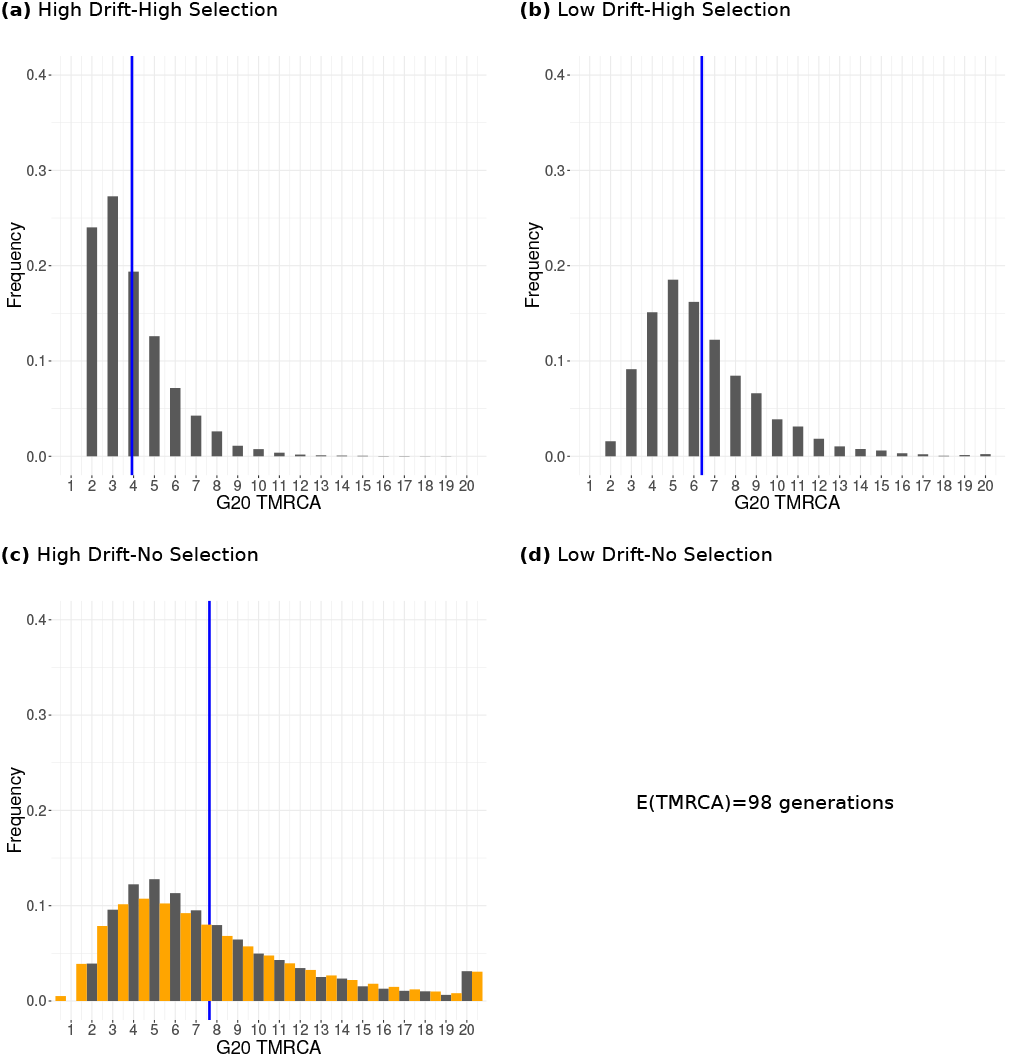
Frequency distribution of the Time to the Most Common Ancestor of progenitors constituting the last simulated generation. *G*_20_ TMRCA distribution (in grey) was obtained under HDHS (a), LDHS (b), HDNS (c) with mean TMRCA indicated as a blue vertical line. In (c), we plotted in gold the theoretical expectation of TMRCA distribution following Eq: (11). Note that under LDNS, theoretical expectations for TMRCA reached 98 generations, while our simulations were run for 20 generations. We therefore discarded the corresponding graph.

We next assessed the impact of selection on *N_e_* and compared different estimates, either based on TMRCA (Eq: (9)), or on the variance in offspring number (Eq: (10)), or on the cumulated response to selection (Eq: (8)). Values obtained are summarized in Tab. 3c. We found that in the absence of selection, *N_e_* estimated from the mean TMRCA were close to the actual number of reproducing individuals (4.8 for HDNS and *>*10 for LDNS), while they were much smaller under both selection regimes (2.5 for HDHS and 3.3 for LDHS). The observed differences between 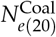 and the harmonic mean of 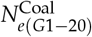 revealed a strong influence of the first generation on the adaptive dynamics. When *N_e_* was computed from the variance in offspring number, estimates without selection (4.1 under HDNS and 42 under LDNS) were close to the actual number of reproducing individuals. Finally, *N_e_* estimates from the cumulated response to selection fell within the same range as the ones from the variance in offspring number in both selection regimes. In summary, most *N_e_* estimates were close to the actual number of reproducing individuals in the absence of selection but high selection intensity strongly reduced *N_e_* estimates.

### Stochasticity in the response to selection

We addressed the qualitative nature of selection response focusing on its linearity. To do so, we measured in each family the average genetic gain per generation over 2000 simulations by fitting a linear regression model. The average genetic gain was 0.49 DTF/generation under HDHS, and 1.1 DTF/generation under LDHS (Tab. 3c). Associated *R*^2^ *>* 0.95 indicated an accurate fit of the data to the linear model. Yet, large standard deviations around these estimates (0.2 and 0.27 for HDHS and LDHS, respectively) pointed either to high stochasticity or a non-linear response. Single simulations indicated non-linear response (Fig. S3). Noteworthy, a strong response was observed between *G*_0_ and *G*_1_ (*G*_0_ *G*_1_ Fig. S3) with similar values in HDHS and LDHS, around 1.6 DTF/generation (Tab. 3c). Subsequently, simulations displayed discontinuities with abrupt changes of slopes at some generations, a signal compatible with the fixation of new mutations (Fig. S3). In order to characterize such discontinuities, we fitted a linear segmentation regression on individual simulations from *G*_1_ and onwards. We estimated the number of breakpoints (i.e. slope changes), the corresponding slopes, and the first and greatest slope based on AIC minimization (Durand *et al.* 2010). The first slope described an average gain of 0.59 DTF/generation in the HDHS regime, and almost twice (0.96 DTF/generation) in the LDHS regime (Tab. 3c). These values were lower than those observed in *G_0_ G_1_*.

Those results are consistent with a *G*_0_*G*_1_ response resulting from the recruitment of initial genetic variance, independently of the population size, and a later response based on mutational variance being less effective in small than in large populations. To confirm those results, we performed a principal component analysis (PCA) and explored correlations between input parameters: initial additive genetic variance 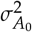, mutational variance 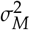 and residual variance 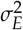, and descriptors of the response to selection : *G*_0_*G*_1_ response, number of breakpoints, first slope and greatest slope. In line with our interpretation, irrespective of the selection regime, 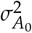 positively correlated with *G*_0_ *G*_1_, and 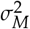 positively correlated with the first (after *G*_1_) and greatest slope (Fig. S4). Note that this stochastic process of mutation occurrence and fixation resulted in large differences among replicates, as illustrated by the breadth of the response (shaded areas in Fig. 1).

### Evolution of genetic diversity

Because of the well-established role of standing variation in selection response, we focused on its temporal dynamics. Standing variation in our experiment consisted in residual heterozygosity found in the initial inbred lines. Starting with a mean residual heterozygosity of 3.0 × 10^−3^ at *G*_0_ (Tab. 3c), we observed a consistent decrease throughout selfing generations until the mutation-drift-equilibrium was reached (Fig. S5). The mean values reached ≈7.0 × 10^−4^ at *G*_20_ without selection, and ≈8.0 × 10^−4^ with selection (Tab. 3c) irrespective of the census population size.

Concerning the number of polymorphic loci, a mutation-drift-equilibrium was reached in all cases except for the LDNS selection regime (Fig. S6). The equilibrium value depended on the census population size: around 6 polymorphic loci with high drift (HDHS and HDNS), 40 polymorphic loci under LDHS, and *>* 66 polymorphic loci after 20 generations under LDNS (Tab. 3c (c) and Fig. S6). Altogether, our results show that the mean heterozygosity was affected neither by drift, nor by selection, but instead by the mutation rate. On the contrary, the number of polymorphic loci depended on the census population size.

### The dynamics of *de novo* mutations

Evolution of frequencies of new mutations revealed three fates: fixation, loss, and rare replacement by *de novo* mutation at the same locus. The four regimes strikingly differed in their mutational dynamics (Fig. 3). Under HDHS, most mutations quickly reached fixation (3.8 generations), with an average of 7.7 fixed mutations/population in 20 generations (Tab. 3c). The corresponding Low Drift regime (LDHS) displayed longer fixation time 5.9 generations, and an average of 10 fixed mutations/family (Tab. 3c). Regimes without selection tended to exhibit a depleted number of fixed mutations, with no fixation under LDNS after 20 generations. Variation around the mean fixation time was substantial across all regimes (Fig. S7). In sum, HDHS was characterized by the fast fixation of new mutations whose direction corresponded to the direction of selection: 53% were fixed within 2 to 3 generations which contrasted to 15% under LDHS or 17% under HDNS. Selection therefore increased the number of fixed mutations while decreasing their fixation time.

**Figure 3.**
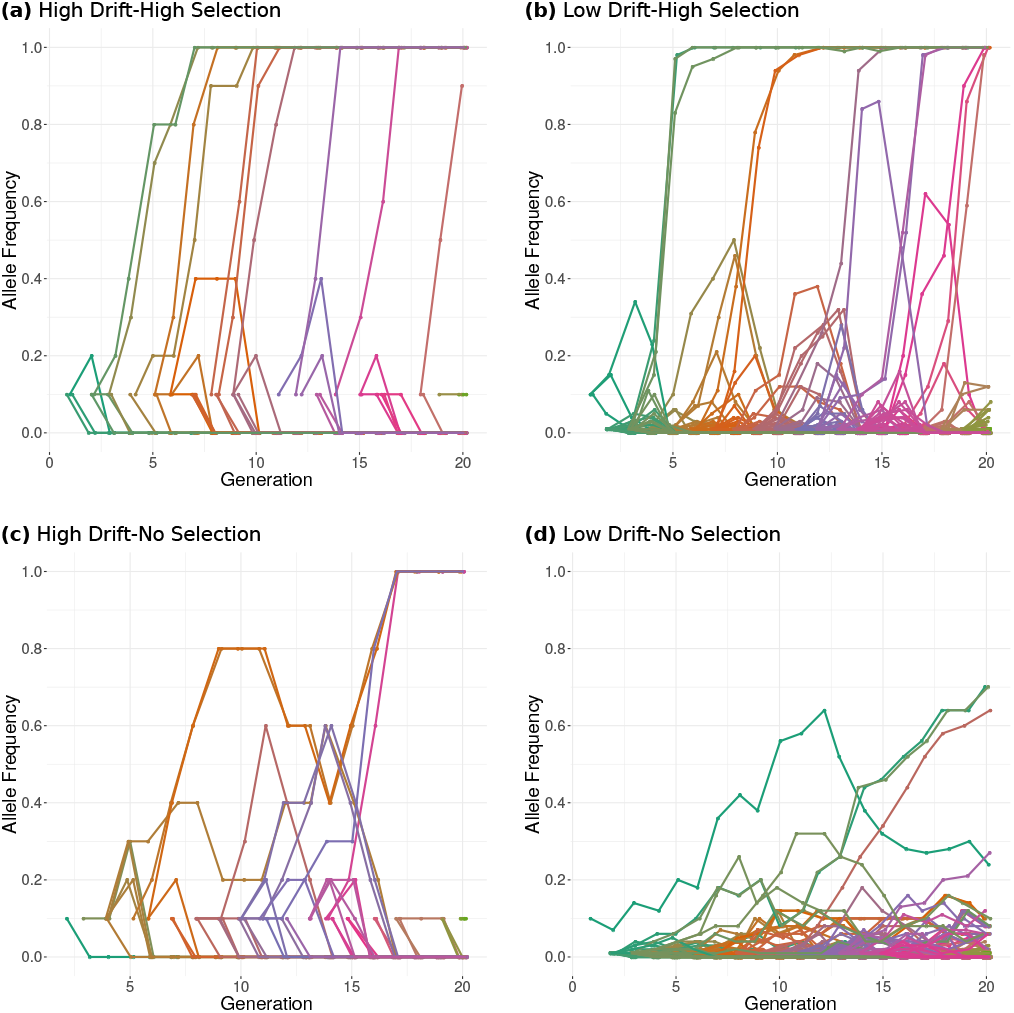
Evolution of allele frequencies within families under four simulated regimes. Examples of mutational fates are given for HDHS (a), LDHS (b), HDNS (c), LDNS (d). Mutations are recorded only when occurring in one of the selected progenitors, and corresponding frequencies are computed over all selected individuals. For example under High Drift regimes, the initial frequency of a mutation occurring in any given progenitor within a family is 1 ÷ (2 × 5) as 5 diploid individuals are selected at each generation. Under Lower Drift regimes, the mutation initial frequency equals 1 ÷ (2 × 50).

### Effects of mutations

Beyond fixation time, a key aspect of our work was to investigate the impact of drift and selection on the type of fixed mutations, best summarized by their genotypic effects. In order to do so, we compared the distribution of incoming *de novo* mutations to that of fixed mutations. We observed a strong depletion of deleterious mutations together with a striking enrichment in beneficial mutations under the two Selection regimes, HDHS (quantile 5%=−0.02, median value=0.43, 95% quantile=1.3) and LDHS (quantile 5%=0.011, median value=0.66, 95% quantile=1.7) (Fig: 4). We also derived a theoretical expectation from Kimura’s allele fixation probability using the selection coefficient computed in the case of truncation selection (Eq: 16). Accounting for the specificities of our selection procedure we found under both selection regimes, a slight excess of detrimental mutations, and a large excess of beneficial mutations as compared to Kimura’s predictions. Note however that, comparatively, the excess of detrimental mutations in the simulations compared to theoretical expectations was more reduced under HDHS than under LDHS (Fig: 4).

**Figure 4.**
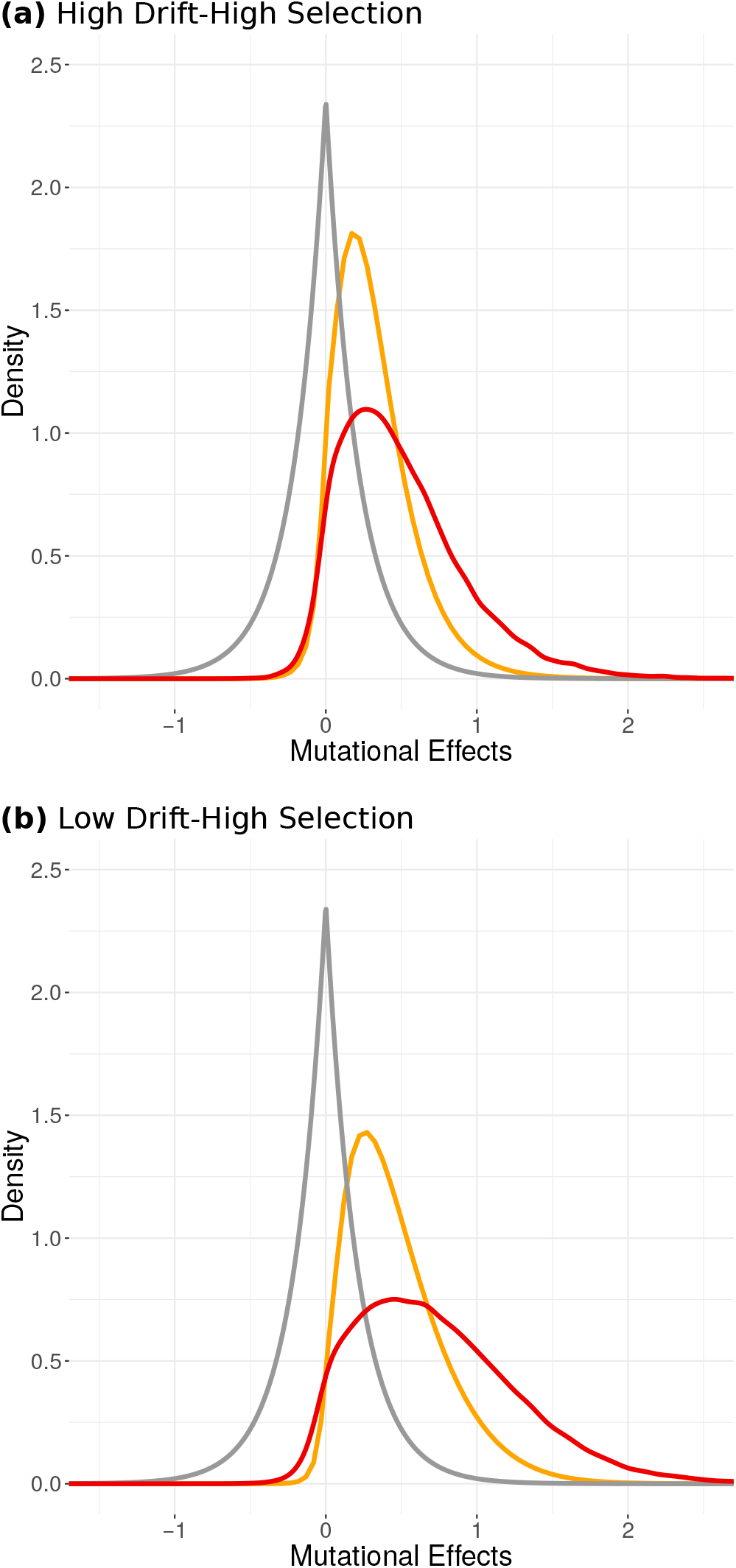
Distribution of effects of *de novo* and fixed mutations under High Selection intensity regimes. Density distributions for the HDHS (a) and the LDHS (b) regime are shown for all *de novo* mutational effects in grey — reflected exponential distribution —, and fixed mutations over 2000 simulations in red. Theoretical expectations from (Eq: 16) are plotted in gold.

As expected, selection generated a relation between the average size of a mutation and its time to fixation : the higher the effect of the mutation, the lower the time to fixation (Fig. 5 (a) and Fig. 5 (b)). Comparison between HDHS and LDHS revealed interesting features: under high drift, the average effect of mutations fixed was lower and variance around mutational effects tended to decrease correlatively with fixation time so that large size mutations were all fixed during the first generation while they persisted at subsequent generations under Low Drift (Fig. 5 (a) and Fig. 5 (b)).

**Figure 5.**
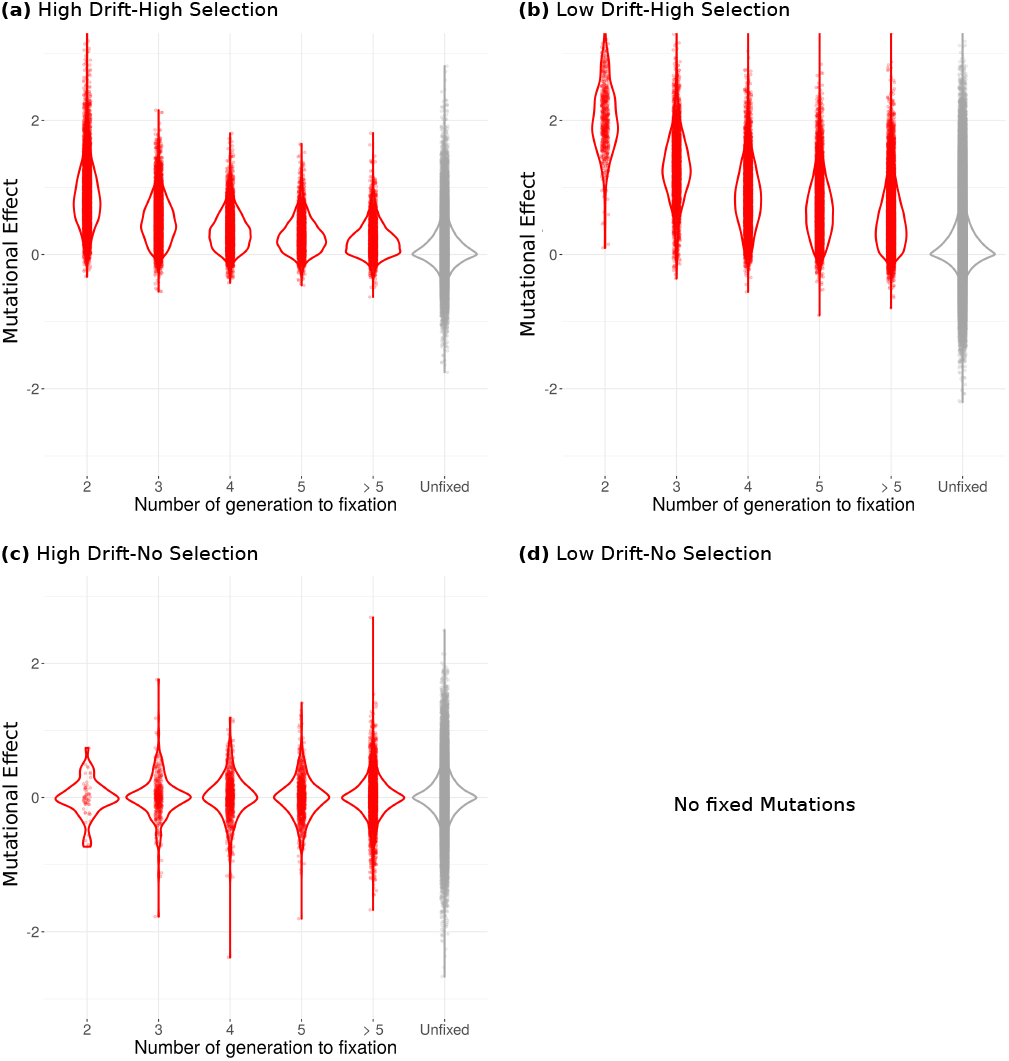
Violin plots of raw mutational effects according to fixation time under three simulated regimes. Plots are indicated for fixed (red) and lost (grey) mutations under HDHS (a), LDHS (b) and HDNS (c). Note that under LDNS, we obtained very few fixed mutations so that we were unable to draw the corresponding distribution.

In sum, our two selection regimes lead to an enrichment of beneficial mutations. Compared with LDHS, HDHS regime fixed fewer detrimental mutations but the average effect of fixed beneficial mutations was smaller.

### Covariation between mutational and environmental effects

A puzzling observation was that normalizing raw mutational effects by the environmental standard deviation of selected individuals translated into a distortion of the distribution so that the median value of fixed effects increased by 0.29 (from 0.43 to 0.72) under HDHS and by 0.2 under LDHS (Tab. 3 and Fig. S8). Similarly, 95% quantile increased by 1.2 (from 1.3 to 2.5) under HDHS and 0.66 (from 1.7 to 2.4) under LDHS. Hence, normalization distortion resulted in much more similar fixed mutations effects distribution under HDHS and HDNS. This was due to a non-zero negative genetic-environment covariance in selected individuals. Indeed, conditioning on the subset of selected individual, we obtained negative estimate of Cov(*G* _|selected_, *E* _|selected_) both under HDHS and LDHS, with a median value (respectively 5% and 95% quantile) of −0.11 (respectively −0.88 and 0.029) under HDHS and −0.37 (respectively −1.1 and −0.072) under LDHS. In contrast, with no selection, values of randomly chosen individual Cov(*G*_|random_, *E*_|random_) were centered around 0 as expected. The evolution of Cov(*G* _|selected_, *E* _|selected_) through time (Fig. S9) evidenced a high stochasticity among generations but no temporal autocorrelation (Fig. S9). In other words, because of the negative correlation between residual environmental effects and genetic effects induced by selection, mutational effects tightly depended on their environment of selection.

## Discussion

Population and quantitative genetics provide theoretical frameworks to investigate selection responses and underlying multilocus adaptive dynamics. Here, we focused on Saclay DSEs which were specifically designed to depict the evolutionary mechanisms behind the response to selection of a highly complex trait — with a high mutational target — in small populations evolving under truncation selection (1% of selected individual), limited recombination (complete selfing) and limited standing variation.

Our main motivation was to explore how such a combination of unusual conditions, at the limits of parameters boundaries of classic models, can sustain the long-term maintenance of additive genetic variation and a significant selection response with no observed load (annual field observations). In this purpose we (1) devised forward individual-based simulations that explicitly modeled our Saclay DSEs and were calibrated on observed values of initial, mutational and environmental variances, and (2) relied on theoretical predictions to investigate the interplay of evolutionary forces and patterns associated with fixation of mutations.

### The broadness of the mutational target sustains long-term mutational input

The three determinants of the observed selection response were best summarized by three variance components namely, the initial additive variance 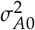, the mutational variance 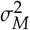, and the environmental variance 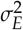 (Fig: S4). Quantitatively, we demonstrated the importance of both initial standing variation and the necessity of a constant mutational input to explain the significant selection response in the two Saclay DSEs (Fig: 1 & S4). This result was consistent with previous reports (Durand *et al.* 2010, 2015) and showed that the first selection response between *G*_0_ and *G*_1_ was correlated with 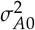, while response in subsequent generations was mainly determined by 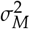 (Fig: S4). In our simulations, we chose initial values for variance components that closely matched previous estimates in the Saclay DSE derived from the MBS inbred line (Durand *et al.* 2010). The small value for initial additive variance came from the use of commercial inbred lines in our experimental evolution setting. It sharply contrasted with more traditional settings where distant genetic material and crosses are often performed to form an initial panmictic population on which selection is applied (Kawecki *et al.* 2012). While crucial in the first generation (Fig: S4), 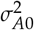 was quickly exhausted. Hence, we showed that the long-term selection response was sustained by a strong mutational variance. The chosen variance 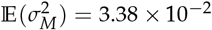, corresponded to an expected mutational heritability of 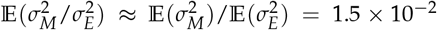 (in units of residual variance per generation). This value, observed in our setting for flowering time, stands as a higher bound to what was previously described in other species/complex traits (Keightley 2010; Walsh and Lynch 2018).

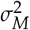 is intimately linked to the broadness of the mutational target, a key parameter of our setting. While decreasing the genomic mutation rate (U) - either by modifying the number of loci or by the per-base mutation rate - we observed at first sight a stronger average response to selection (Fig. S10). This is because incoming *de novo* mutational effects increase correlatively with decreasing U, to maintain 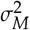 constant. Note that stochasticity of the response is boosted by scarcity of strong effect mutations that are preferentially fixed (Fig. S10). If instead of conditioning on 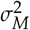, we conditioned on the distribution of incoming *de novo* mutational effects, responses under low values of U were drastically reduced (Fig. S10). These results further highlight that adaptive evolution results from a subtle balance between mutation, drift and selection.

We implemented an additive incremental mutation model (Clayton and Robertson 1955; Kimura 1965; Walsh and Lynch 2018). This model assumed non-limiting mutational inputs, and has been shown to be particularly relevant in systems where, just like ours, effective recombination is limited (Charlesworth 1993; Walsh and Lynch 2018). Alternative model such as the House Of Cards (HoC) that sets random allelic effect upon occurrence of a new allele (Kingman 1978; Turelli 1984) — rather than adding effects incrementally — would have likely resulted in smaller estimate of 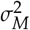 (Hodgins-Davis *et al.* 2015). Whether the incremental model or the HoC or a combination of both such as the regression mutation model (Zeng and Cockerham 1993) was better suited to mimic our Saclay DSEs is an open question. However several lines of evidence argue in favour of a nonlimiting mutational input in our setting. First, the architecture of maize flowering time is dominated by a myriad of QTLs of small additive effects (Buckler *et al.* 2009). Over 100 QTLs have been detected across maize lines (Buckler *et al.* 2009), and over 1000 genes have been shown to be involved in its control in a diverse set of landraces (Romero Navarro *et al.* 2017). Second, in Saclay DSEs alone, transcriptomic analysis of apical meristem tissues has detected 2,451 genes putatively involved in the response to selection between early and late genotypes, some of which being interconnected within the complex gene network that determines the timing of floral transition (Tenaillon *et al.* 2018). This suggests that not only the number of loci is considerable, but also that their connection within a network further enhances the number of genetic combinations, and in turn, the associated phenotypic landscape.

The breadth of the mutational target is a key parameter for adaptation (Höllinger *et al.* 2019). Altogether, our results suggest that despite a small number of segregating loci (Tab. 3c and Fig. S6) expected under HDHS, adaptive variation was continuously fueled by a large mutational target which translated into a long-term selection response. In other words, the large mutational target compensates for the small population sizes, and triggers the long-term maintenance of adaptive diversity at the population level after the selection-drift-mutation equilibrium is reached, i.e. after three to five generations. Noteworthy the expected level of heterozygosity in our controls (No Selection models, NS) corresponded to neutral predictions (Crow and Maruyama 1971; Kimura 1969).

### Quick fixation of *de novo* mutations drive Saclay DSEs selection response

The observed fixation time of mutations without selection is expected under standard neutral theory. The Kingman coalescent indeed predicts a TMRCA around 8 generations for a population size of 5 which matched closely our observed value of 7.6 obtained under HDNS. With selection, instead, we observed a quick fixation of mutations in three to four generations under HDHS. Likewise, the number of fixed mutation increased from 2.3 in HDNS to 7.7 in HDHS (Tab. 3). Note that while one would expect emerging patterns of hard sweeps following such rapid mutation fixation, our selfing regime which translated into small effective recombination likely limits considerably genetic hitchhiking footprints, so that such patterns may be hardly detectable.

Short fixation times made the estimate of effective population sizes challenging. We used two estimates of *N_e_* to shed light on different processes entailed in HDHS stochastic regime. These estimates were based on expected TMRCA and on the variance in the number of offspring (Crow and Kimura 1971), respectively. We found the latter to be greater than the former. This can be explained by the fact that selection is known to substantially decrease effective population size on quantitative trait submitted to continuous selection, because of the increase in covariance between individuals due to selection (Santiago and Caballero 1995), and because selection on the phenotypic value acts in parts on non-heritable variance (*i.e.*, on the environmental variance component of *V_P_* (Chantepie and Chevin 2020)). Note that our estimates of *N_e_* — based on coalescence times computed from the known genealogical structure — allow to account for these effects. However, this is not without drawback: TMRCA estimates were much shorter than expected, a result consistent with the occurrence of multiple merging along pedigrees, *i.e.* multiple individuals coalescing into a single progenitor. In fact, multiple merger coalescence models may be better suited to describe rapid adaptation than the Kingman coalescent (Neher 2013).

Both fixation time and probability depend on the selection coefficient *s* and the initial frequency of the mutation in the population. In our setting, conditioning on its appearance in the subset of selected individuals, the initial frequency of a mutation was 0.10, which was unusually high and translated into selection and drift exerting greater control over mutations. In contrast, in more traditional drift regimes, even when an allele is strongly selected (2*N_e_s* ≫ 1), drift dominates at mutation occurrence, *i.e.* with two absorbing states for allele frequency near zero and one (Walsh and Lynch 2018). Under HDHS regime, selection induces repeated population bottlenecks that change the structure of the pedigrees and translate into a decrease in effective population size compared to the HDNS regime. Drift and selection can therefore not be decoupled, and they do not act additively.

### High stochasticity promotes the fixation of small effect mutations

Interplay between drift and selection promoted stochasticity in our setting, which manifested itself in various ways : (i) through the selection response, with different families exhibiting contrasting behaviors, some responding very strongly and others not (Fig. 1); (ii) through the dynamics of allele fixation (Fig. 2 & 3); and (iii) through the distribution of Cov(*GE*) (Fig. 6). Stochasticity tightly depends on census population size (Hill 1982a,b). Unexpectedly, however, we found a benefice to stochasticity as illustrated by a bias towards the fixation of advantageous mutations compared with the expectation (Fig. 4). Comparison of the distributions of the mutational raw effects indicated that, among advantageous mutations, a greater proportion of those with small effects were fixed under the High Drift than under the Low Drift regime (Fig. 4 (a) versus (b)). This result echoes those of Silander *et al.* (2007), who showed — using experimental evolution with bacteriophage — that fitness declines down to a plateau in populations where drift overpowers selection. The authors note: “If all mutations were of small effect, they should be immune to selection in small populations. This was not observed; both deleterious and beneficial mutations were subject to selective forces, even in the smallest of the populations.”.

**Figure 6.**
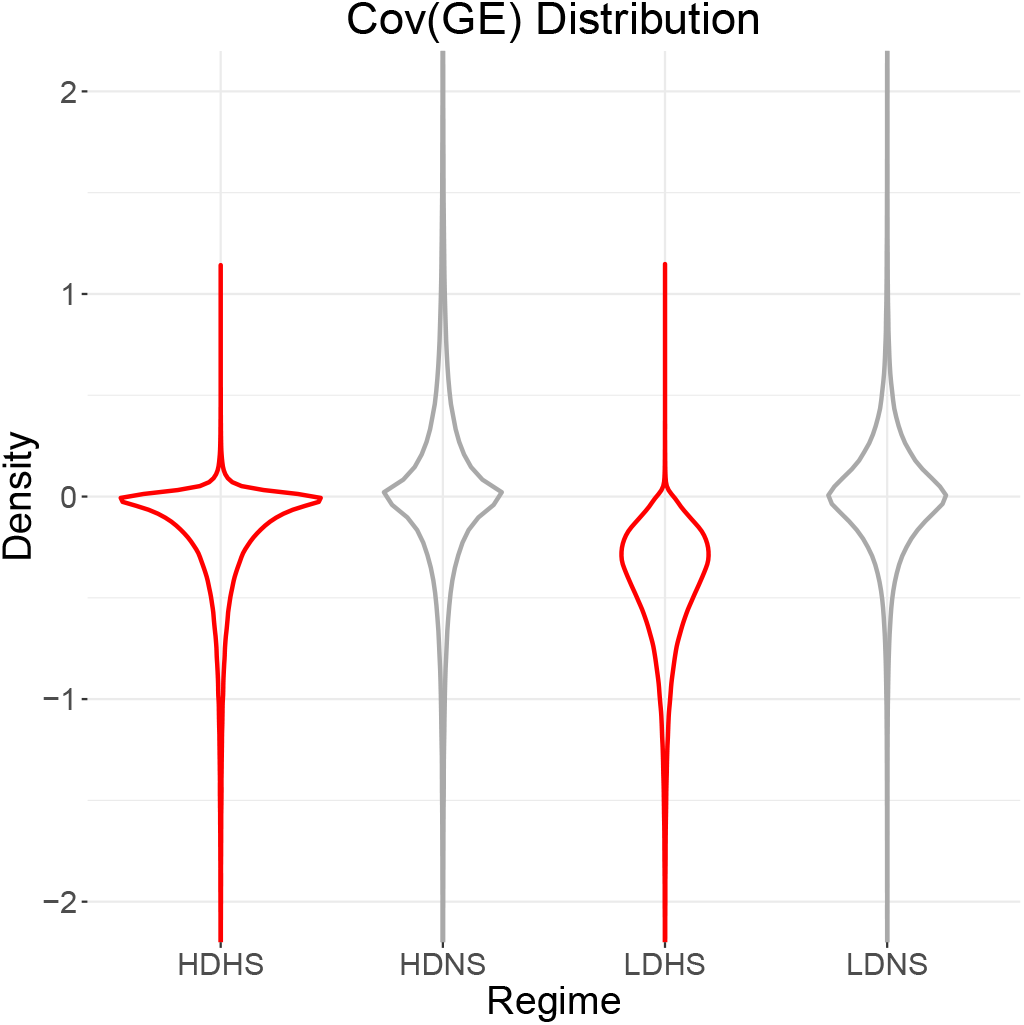
Violin plots of Cov(*G* _|selected_, *E* _|selected_) under four simulated regimes. Violin plots were computed over 2000 simulations and 4 families, four families and across all generations under regimes with High Selection intensity in red (HDHS, LDHS), and regimes with No Selection in grey (HDNS, LDNS).

What are the underlying mechanisms behind this fixation bias? We found a negative covariance between selected genotypes and their corresponding micro-environmental values, that modified the mutational effect to an apparent mutational effect perceived in the particular micro-environment. The negative Cov(*GE*) arose mechanically from selection of two independent random variables, whatever the sampling size as illustrated in Fig. S11 and Fig. S12. This effect evokes the so-called Bulmer effect (Bulmer 1971), that causes a reduction of genetic variance due to the effect of selection on the covariance between unlinked loci. Interestingly, under the High Drift regime, we observed a less negative Cov(*GE*) on average than with a 10 times higher census size (Low Drift). This translated, after dividing by the environmental standard deviation of selected individuals, to a greater apparent effect of small mutations under the High Drift regime. In other words, High Drift-High Selection intensity tends to magnify mutational effects from an environmental perspective. In support of this explanation, normalization by the environmental standard deviation actually erased the difference between the two distributions of mutational effect (under low and High Drift, Fig. S8). Unlike the Bulmer effect however, this one was restricted to the generation of mutation occurrence, but favored long-term fixation of slightly advantageous mutations by a transient increase of their frequency. Because of a significant variance of Cov(*GE*), this effect on small effect mutation fixation was mostly stochastic. Therefore, we interpreted the fixation of a high proportion of slightly beneficial mutations, and their significant contribution to selection response, by the less efficient exploration of the initial distribution per simulation (increasing their prevalence) but the stochastic “help” of a lesser negative Cov(*GE*).

Just like Cov(*GE*), epistatic interactions may further exacerbate stochasticity Dillmann and Foulley (1998). While we showed that *de novo* mutations are fixed sequentially and therefore rarely interact, additive effects of new mutations tightly depend on the genetic background in which they occur (Plucain *et al.* 2014), and mutational history modulates the amount of additive genetic variance (Hill *et al.* 2008). While we have not accounted for them, epistatic interactions may hence result in a distortion of the distribution of mutation effects. Epistasis can also generate asymmetrical selection responses (*Lynch et al.* 1998; Keightley 1996). Because the two initial inbred lines, F252 and MBS, have been intensively selected for earliness (Camus-Kulandaivelu *et al.* 2006; Rebourg *et al.* 2003), we expect in our setting a diminishing return of mutational effects in the Early populations and, conversely, mutations of high effect in the Late populations (Durand *et al.* 2010). Any new mutant occurring within these ‘early’ genetic backgrounds, can either constrain or accentuate phenotypic effects, depending on direction of selection. Altogether, epistasis could explain some discrepancies between observations and simulations, such as the high stochasticity in Saclay DSEs.

### Deficit of fixation of deleterious mutations suggests a limited cost of selection

As expected, we observed that selection decreased the number of segregating polymorphic loci at equilibrium compared to regimes without selection (Tab. 3). Interestingly however, this effect was reduced for small population size. Under High Drift, selection induced an average loss of a single polymorphism at equilibrium (HDHS vs. HDNS, Tab. 3) while under the Low Drift regime over 20 polymorphisms were lost (LDHS vs. LDNS, Tab. 3). A similar trend was recovered at the mutation fixation level where on average 7.7 mutations were fixed under the High Drift-High Selection intensity and only 10 under Low Drift-High Selection intensity. In other words, the 10-fold population increase did not translate into a corresponding increase in the number of segregating and fixed mutations, as if there was a diminishing cost with decreasing population size. Under High Drift (/Low Drift), at each generation 500 (/5000) offspring of 2 × 1000 loci were produced. Considering a mutation rate per locus of 6000 × 30 × 10^−9^, (*i.e.* (Clark *et al.* 2005)), it translated into 180 mutations events (/1800 mutations events). However most mutations are lost as only mutations occurring in the subset of selected individuals survive. The initial frequency of a mutation in this subset, *i.e* of size 5 or 50, is 1/10 under High Drift and 1/100 under Low Drift. In the former, the interplay between the initial frequency and selection intensity allows a better retention of beneficial mutations of small effect (Fig. 4) than in the latter. Interestingly at equilibrium, we also observed a higher level of residual heterozygosity with selection than without, irrespective of population size, suggesting a small impact of selection in the long-term heterozygosity maintenance. Overall, our High Drift-High Selection intensity regime maintains a small, but sufficient number of polymorphisms for the selection response to be significant.

Our selection response evidenced a deficit of fixation of deleterious mutations and hence a modest genetic load (Fig. 4 and S8). We identified three reasons behind this observation. Firstly, in our design, the selection intensity of 1% was applied on the trait. Hence, in contrast to the infinitesimal model for which a high number of polymorphic loci are expected to individually experience a small selection intensity, selection intensity was “concentrated” here on a restricted number of loci, *i.e.* those for which polymorphisms were segregating. Secondly, we applied truncation selection whose efficiency has been demonstrated (Crow and Kimura 1979). The authors noted: “It is shown, for mutations affecting viability in *Drosophila*, that truncation selection or reasonable departures therefrom can reduce the mutation load greatly. This may be one way to reconcile the very high mutation rate of such genes with a small mutation load.” Thirdly, the lack of interference between selected loci in our selection regime may further diminish the selection cost (Hill and Robertson 1966). Reduced interference in our system is indeed expected from reduced initial diversity and quick fixation of *de novo* mutations. Whether natural selection proceeds through truncation selection or Gaussian selection is still a matter of debate (Crow and Kimura 1979). Measuring the impact of these two types of selection on the genealogical structure of small populations including on the prevalence of multiple merging branches will be of great interest to better predict their fate.

This under-representation of deleterious variants echoes with empirical evidence that in crops, elite lines are impoverished in deleterious variants compared to landraces owing to a recent strong selection for yield increase (Gaut *et al.* 2015). Likewise, no difference in terms of deleterious variant composition was found between sunflower landraces and elite lines (Renaut and Rieseberg 2015). Hence, while the dominant consensus is that the domestication was accompanied by a genetic cost linked to the combined effects of bottlenecks, limited effective recombination reducing selection efficiency, and deleterious allele surfing by rapid population expansion (Moyers *et al.* 2018), recent breeding highlights a different pattern. We argue that our results may help to understand this difference because under High Drift-High Selection intensity, a regime likely prevalent in modern breeding, genetic load is reduced. Moreover, our results may provide useful hints to explain the evolutionary potential of selfing populations located at the range margins. Just like ours, such populations are generally small, display both, inbreeding and reduced standing variation (Pujol and Pannell 2008) and are subjected to environmental and demographic stochasticity.

## Conclusion

In conclusion, our High Drift-High Selection intensity regime with non-limiting mutation highlights an interesting interplay between drift and selection that promotes the quick fixation of adaptive *de novo* mutations fueling a significant but stochastic selection response. Interestingly, such selection response is not impeded by the fixation of deleterious mutations so that adaptation in HDHS proceeds with limited genetic load. Our results provide an explanation for patterns highlighted during recent breeding as well as the high colonization ability of small selfing populations located at species range margins. They also call for a better mathematical description of the multilocus adaptive process sustaining the evolution of small populations under intense selection.

## Acknowledgements

We are grateful to Adrienne Ressayre, Elodie Marchadier, Aurélie Bourgais, Sophie Jouanne, Nathalie Galic, Cyril Bauland, and Philippe Jamin who contributed over the years to the field experiments; and to Hélène de Préval and Clément Brusq for their contribution to the simulations.

## Data availability

The complete R script for performing the simulations as well as the summary data for flowering time (2 DSEs and 20 generations) are available at INRAe dataverse (data.inrae.fr): https://doi.org/10.15454/JQABMJ

## Supplementary material

**Figure S1.**
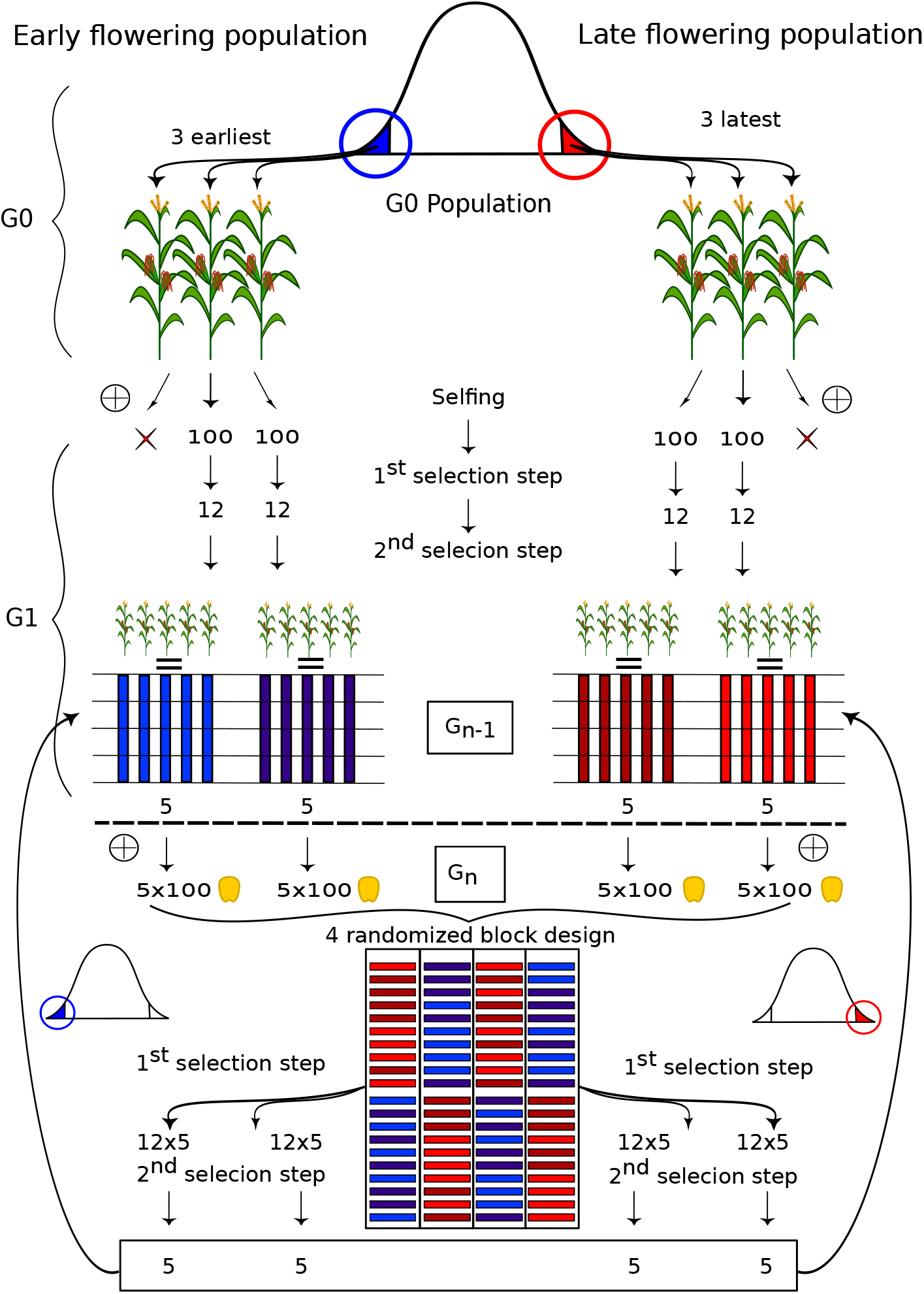
Experimental scheme of Saclay DSEs. For clarity a single scheme is shown but was replicated for the two DSEs. Starting from an inbred *G*_0_ population with little standing variation (*<* 1% residual heterozygosity (Durand *et al.* 2015)), the three earliest/ latest flowering individuals represented in blue/ red were chosen based on their offspring phenotypic values as the founders of two families forming the early/ late population. For the subsequent generations, 10 (≈ 5 per family) extreme progenitors were selected in a two step selection scheme among 1000 plants. More specifically, 100 seeds per progenitor were evaluated in a four randomized-block design, *i.e.* 25 seeds per block in a single row. In a first selection step, the 3 × 4 = 12 earliest/ latest flowering plants among the 100 plants per progenitor were selected in a first step. Then in a second selection step, 10 (≈ 5 per family) individuals were selected within each population based on both flowering time and kernel weight and the additional condition of preserving two progenitors per family from the previous generation.

**Figure S2.**
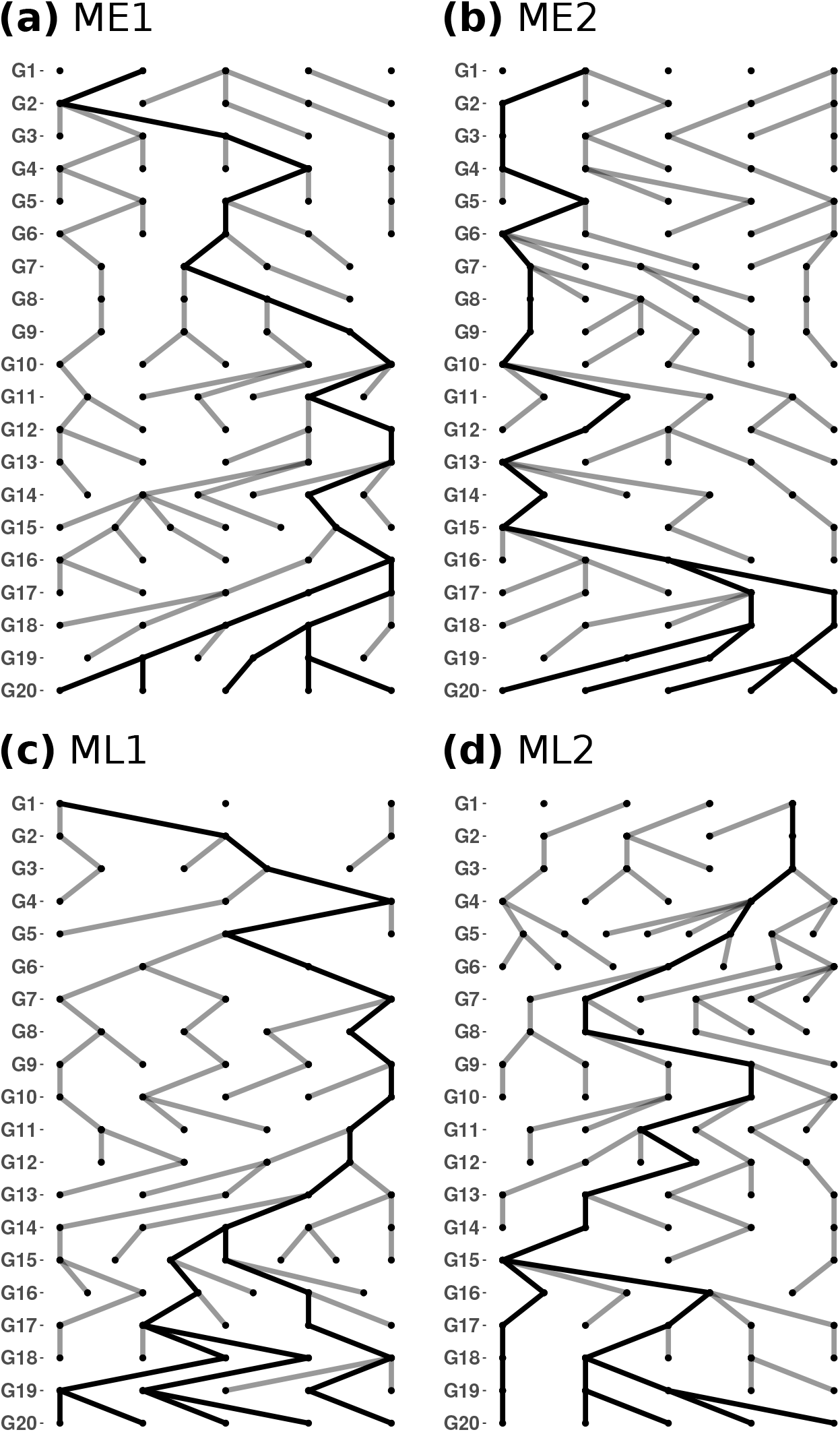
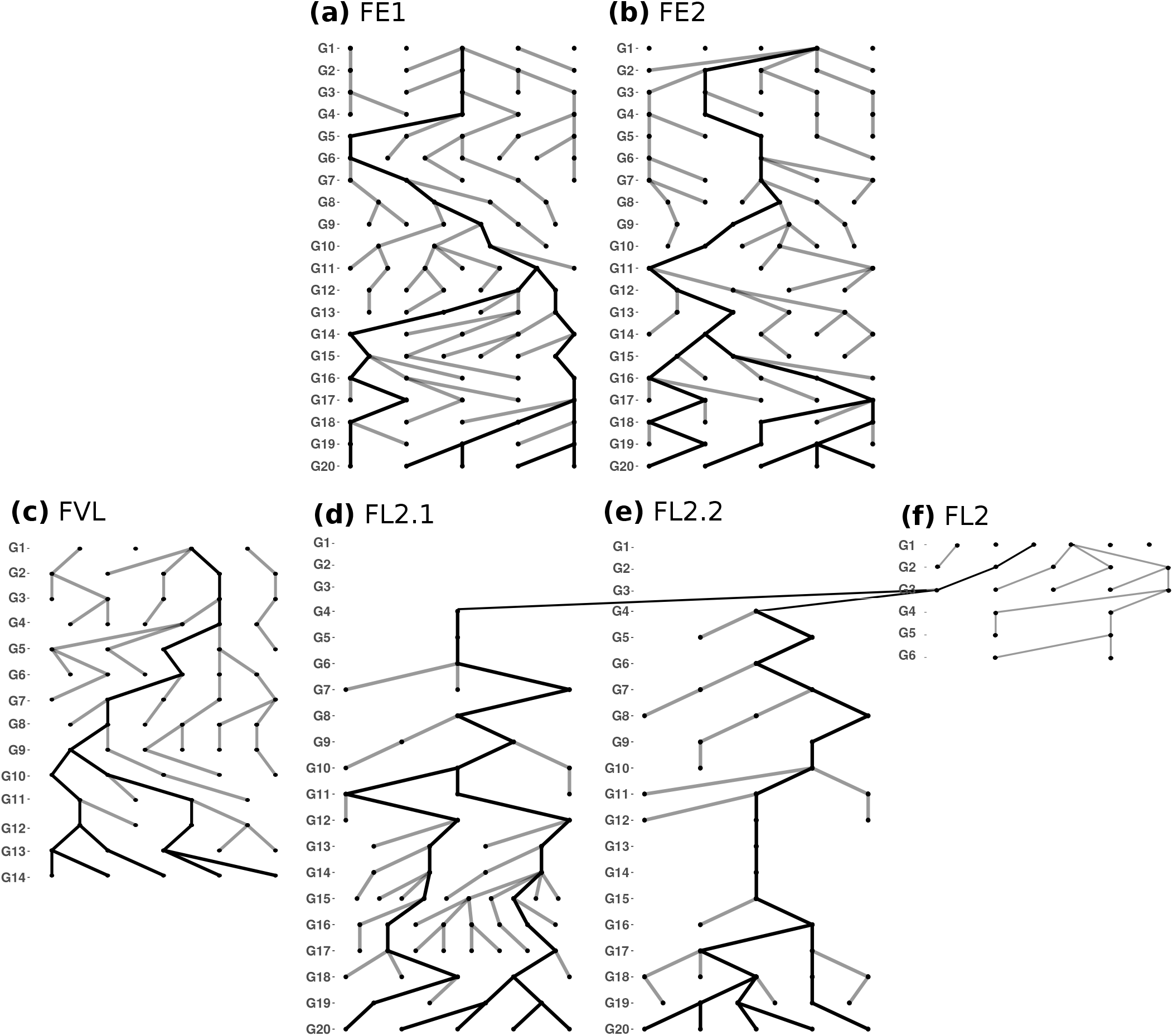
MBS family pedigrees from G1 to *G*_20_. The two early families ME1 (a) and ME2 (b), and the two late families ML1 (c) and ML2 (d) are presented. Each node corresponds to a progenitor selected at a given generation. Each edge corresponds to a filial relationship between a progenitor and its offspring. Thick black lines indicate the ancestral path of the last generation (*G*_20_). **F252 family pedigrees from G1 to *G*_20_.** Two early families FE1 (a), FE2 (b) and two late families FVL (c) & FL2 (f), are represented. FVL (c) could not be maintained after G14 as flowering occurred too late in the season for seed production. Both FL2.1 (d) and FL2.2 (e) were derived from a same individual from FL2 (f) at G3, after FVL was discarded. Each node corresponds to a progenitor selected at a given generation. Each edge corresponds to a filial relationship between a progenitor and its offspring. Thick black lines indicate the ancestral path of the last generation. (*G*_20_)

**Figure S3.**
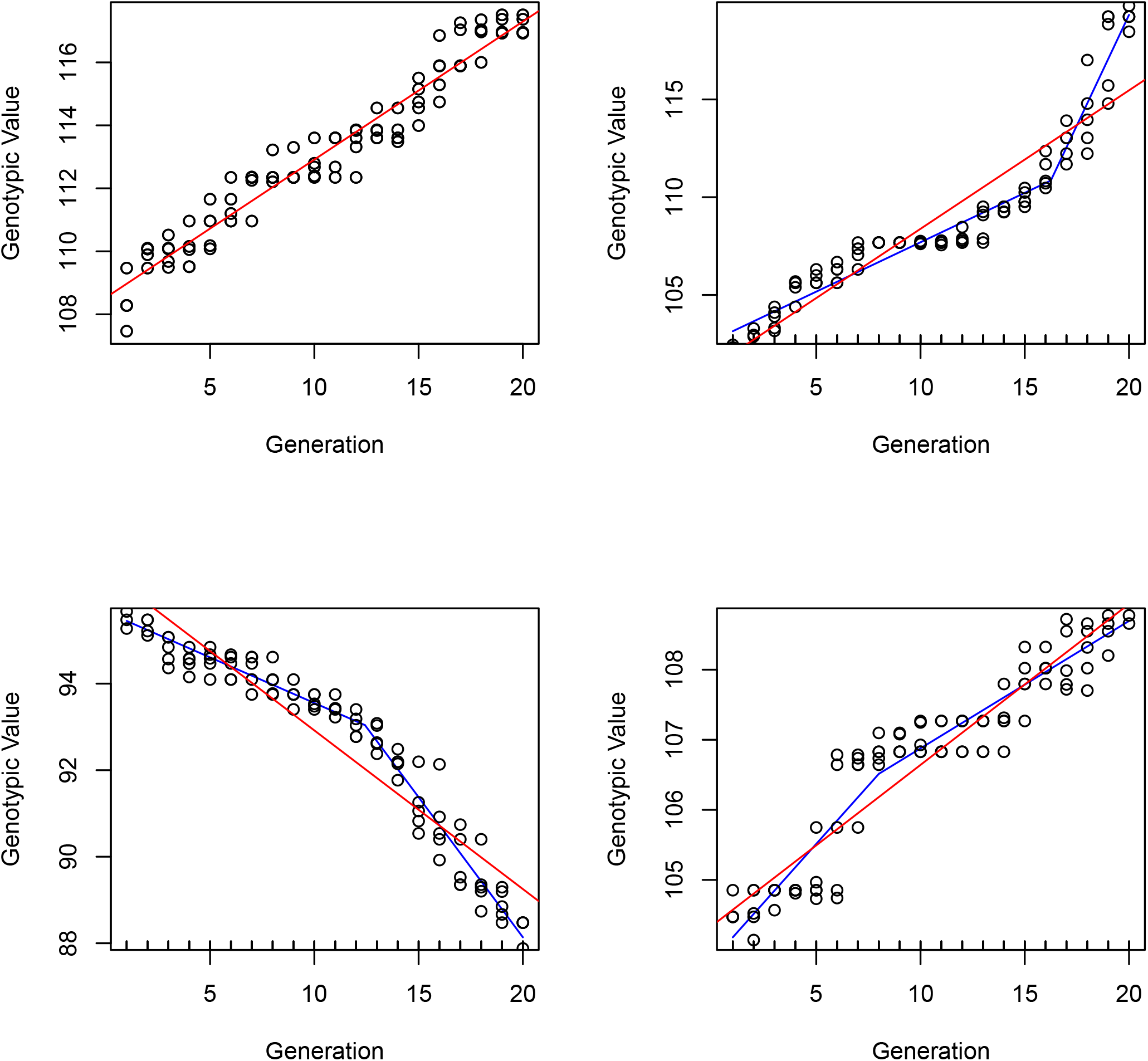
Illustration of simulated non-linear selection response in MBS. Each panel presents the evolution through time (x axis) of the genotypic value (y axis) of the 5 selected individual per family (empty dots). The red lines shows the linear regression of the selected genotypic values through times, while blue lines correspond to the best (AIC criterion) segmented linear model. The top left panel is an example for which a simple linear model fitted best the selection response, while the three others show a diversity of non-linear behaviors.

**Figure S4.**
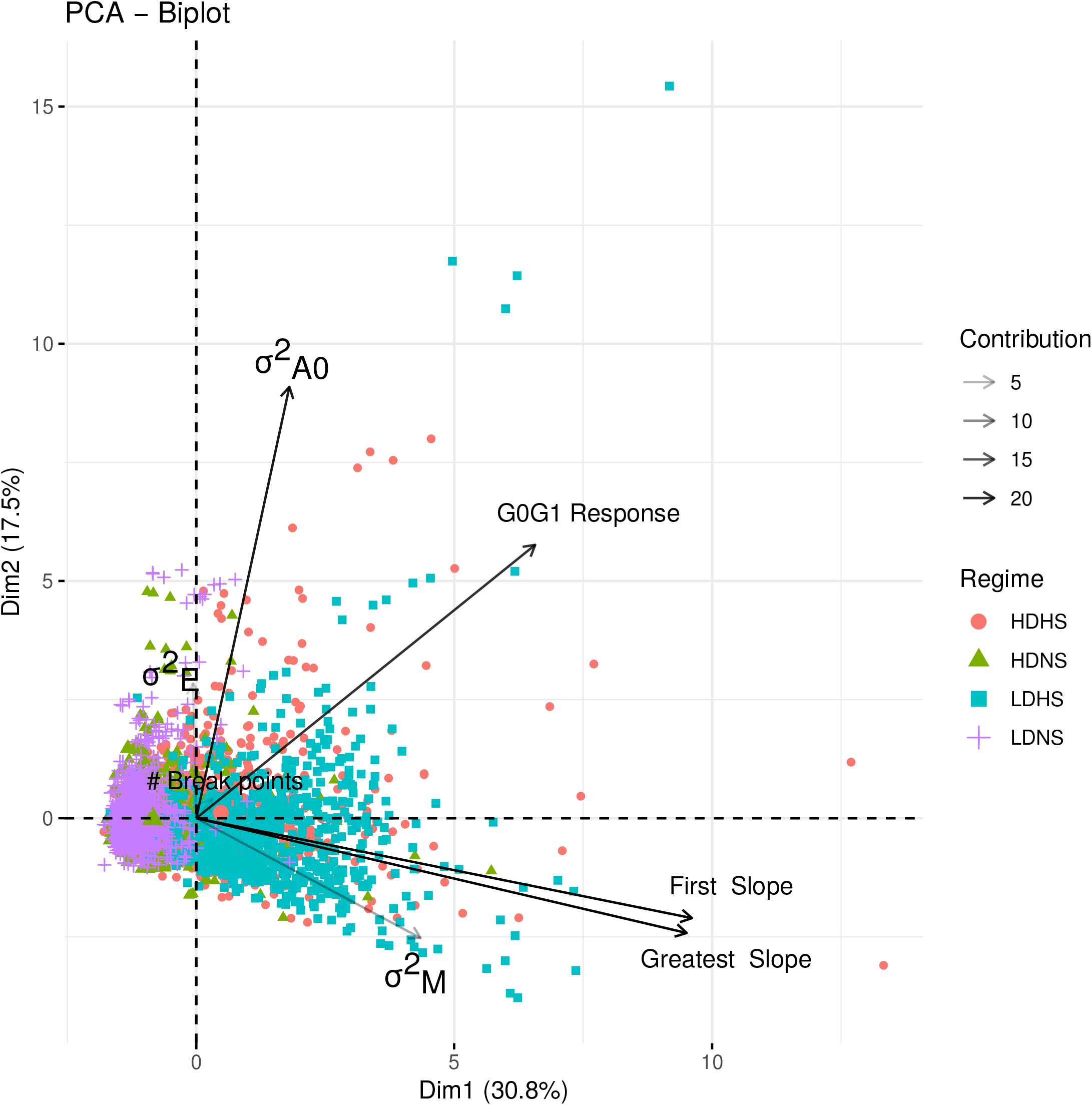
Correlation between model input variables (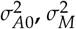 and 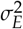) and output variables (*G*_0_*G*_1_ Response, # Breakpoints, First Slope and Greatest Slope). We obtained the output variables by fitting a segmented linear regression to the selection response from *G*_1_ to *G*_20_ in individual. We estimated the number of breakpoints, the corresponding slopes, as well as the first & greatest slope by AIC maximization. In addition we determined the *G*_0_*G*_1_ response. A Principle Component Analysis was carried out on a subset of 200 independent simulations per regime (HDHS, LDHS, HDNS, LDNS). The darker the arrow representing a variable, the higher the intensity of its correlation to the axes.

**Figure S5.**
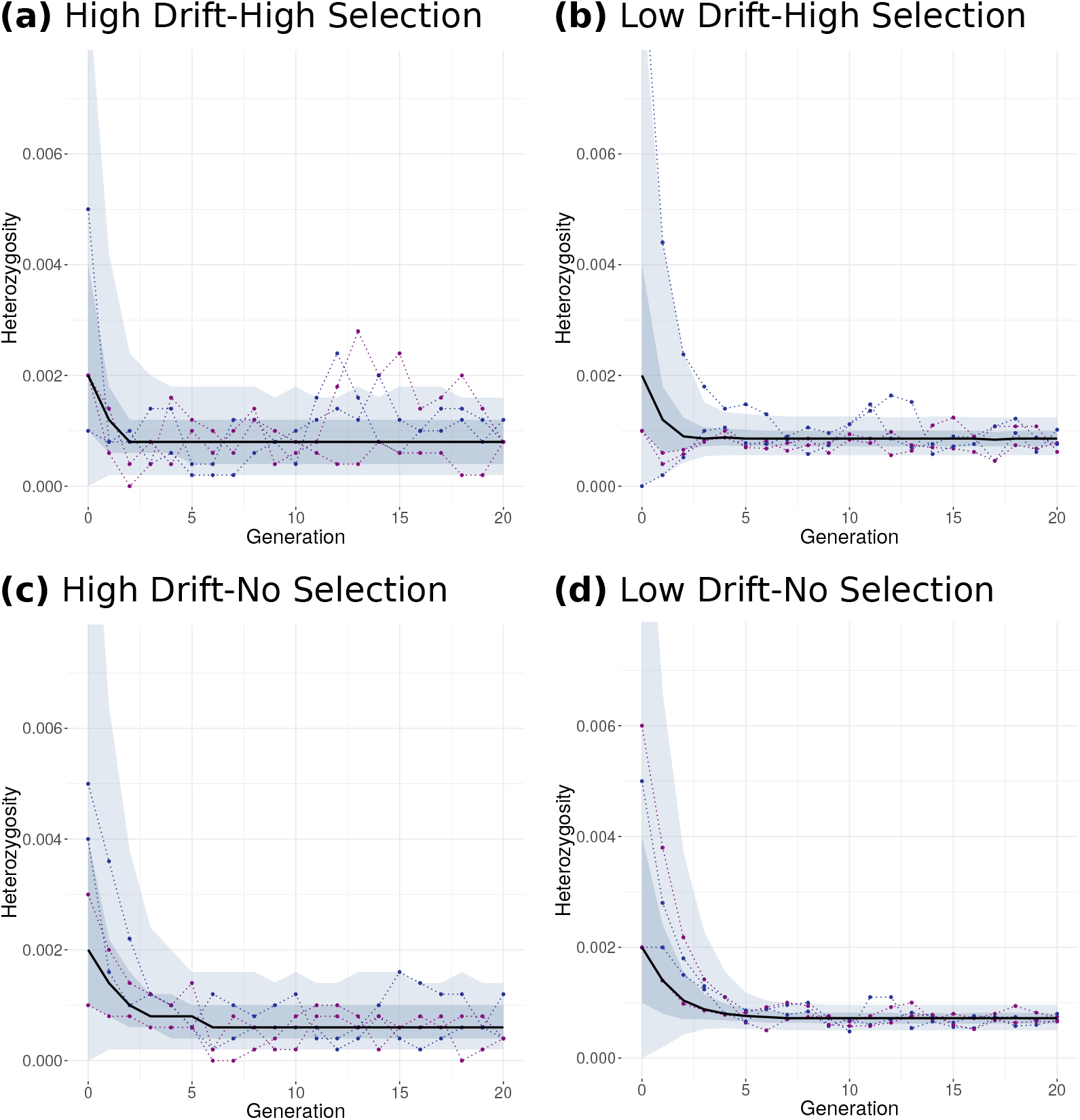
Evolution through time of the per-family mean heterozygosity over all loci, under HDHS (a), LDHS (b), HDNS (c), LDNS (d). The black line represents the median value of the per-family mean heterozygosity. The shaded area corresponds to the 5^*th*^-95^*th*^ percentile (light blue) and to the 25^*th*^-75^*th*^ percentile (dark blue). Four randomly chosen simulated families are represented with dotted line.

**Figure S6.**
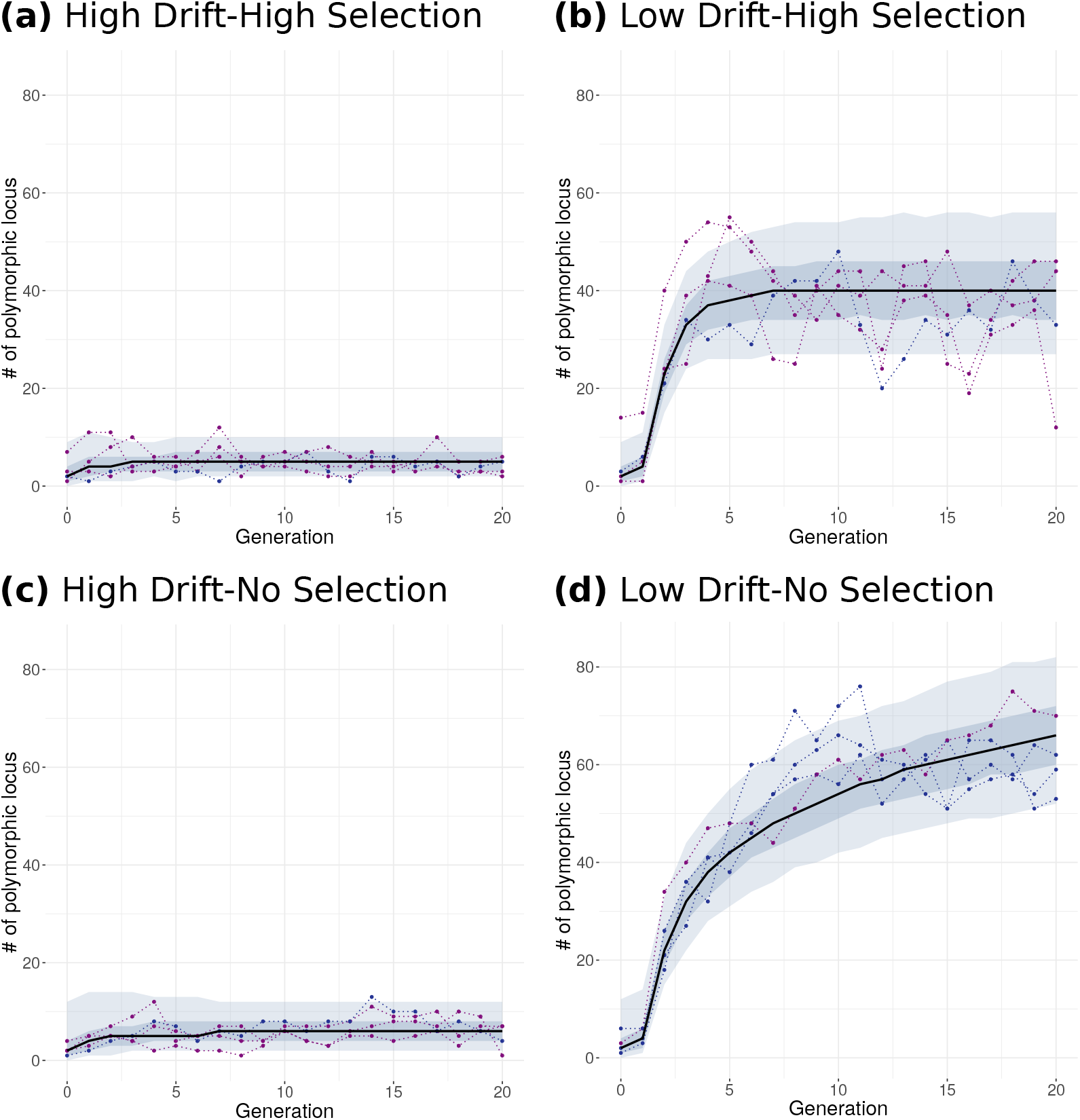
Evolution through time of the per-family mean number of polymorphic loci, under HDHS (a), LDHS (b), HDNS (c), LDNS (d). The black line represents the median value over 2000 simulations. The shaded area corresponds to the 5^*th*^-95^*th*^ percentile (light blue) and to the 25^*th*^-75^*th*^ percentile (dark blue). Four randomly chosen simulated families are represented with dotted line.

**Figure S7.**
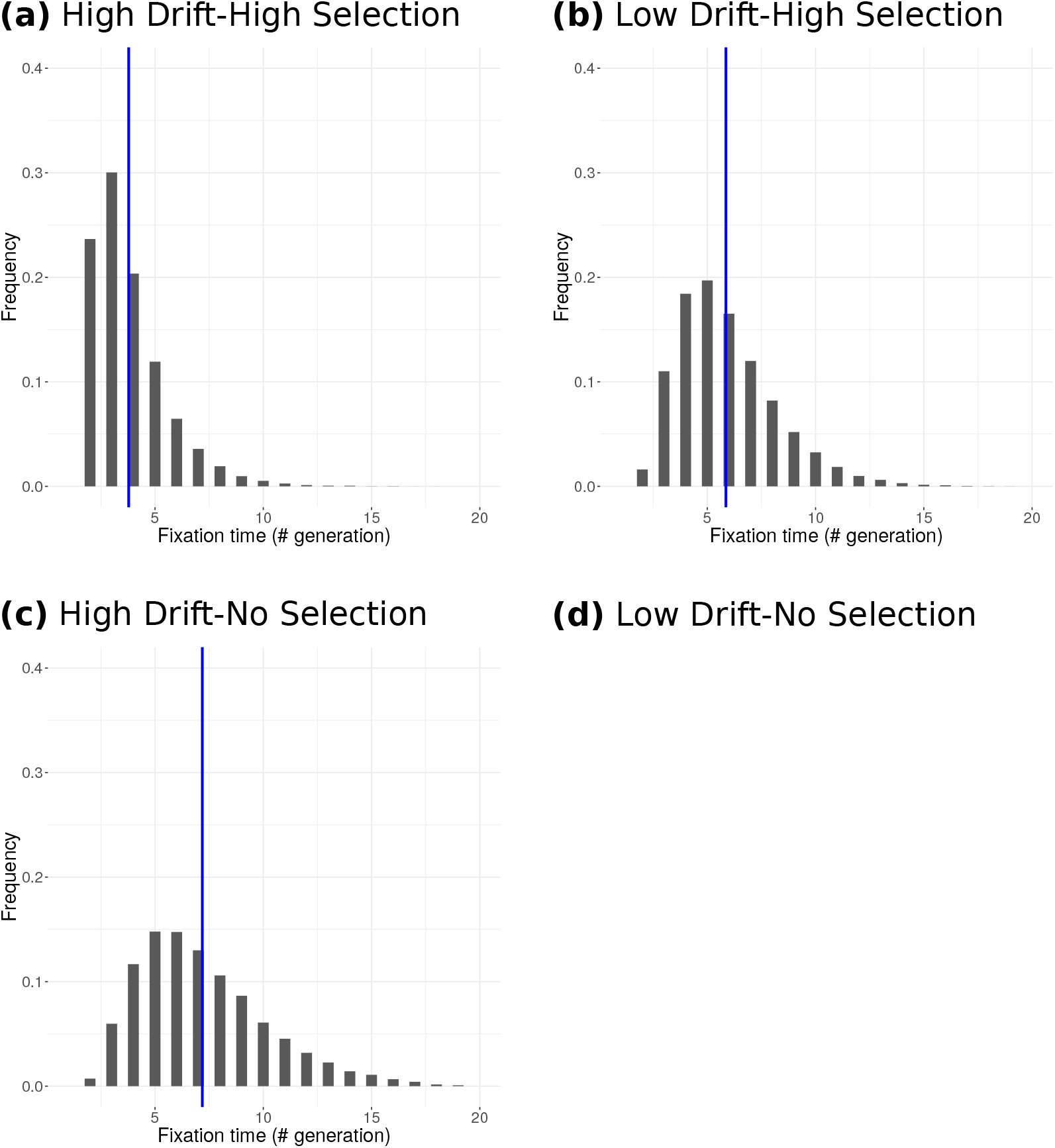
Frequency distribution of mutation fixation times over all simulated families under HDHS (a), LDHS (b), HDNS (c), LDNS (d). Note that under LDNS, we obtained very few fixed mutation so that we were unable to draw the corresponding distribution. Blue vertical lines represent the interpolated median.

**Figure S8.**
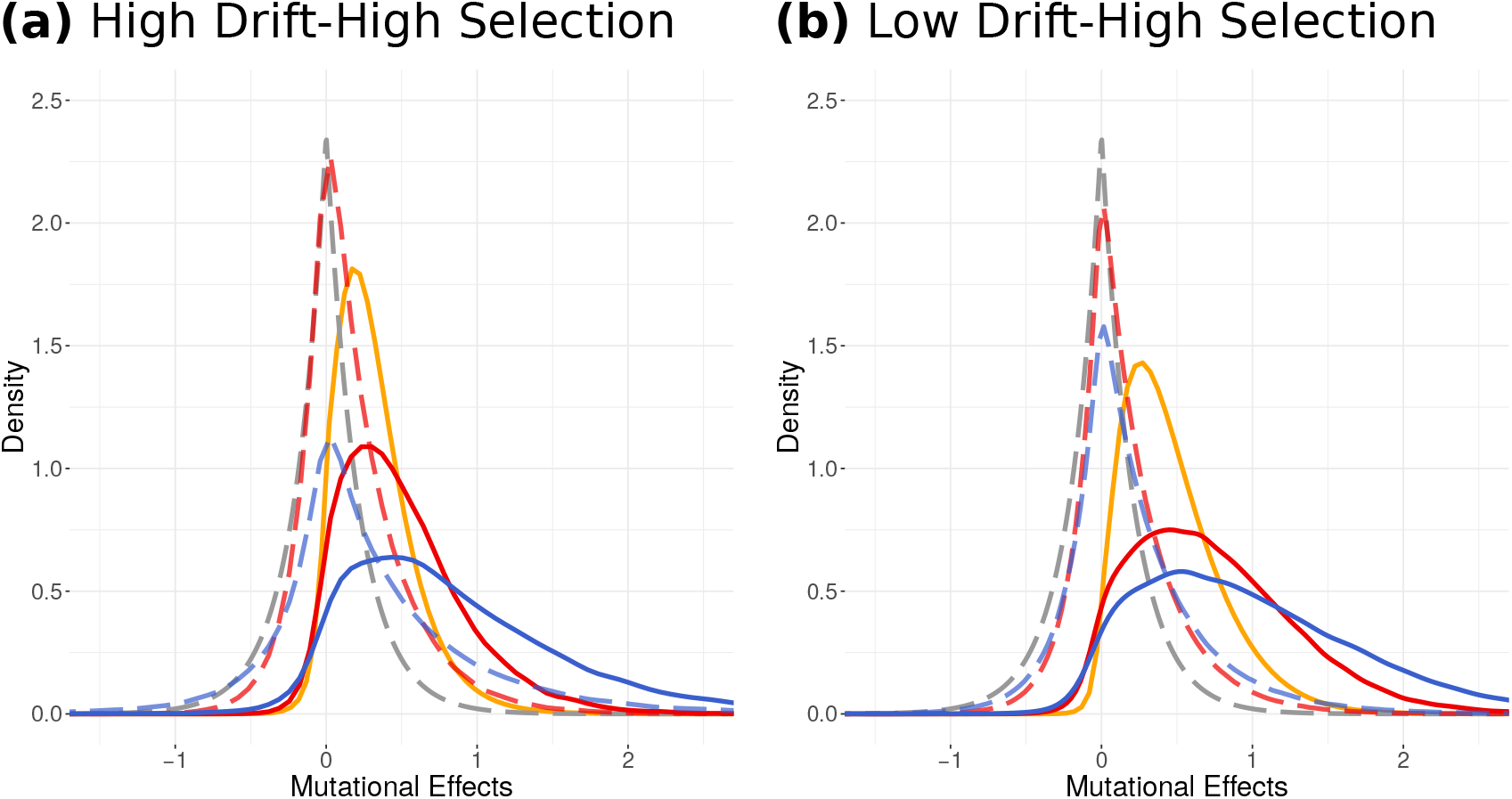
Distribution of mutation effects under HDHS (a), LDHS (b). The dotted lines indicate the distribution of effects (DFE) of incoming *de novo* mutations considering raw effects in all individuals (grey), in selected individuals (red), and effects normalized by environmental variation in selected individuals (blue). The plain lines indicate DFE of fixed mutations following the same colour code. The golden line represents the expected DFE of fixed mutations according to Eq: 16.

**Figure S9.**
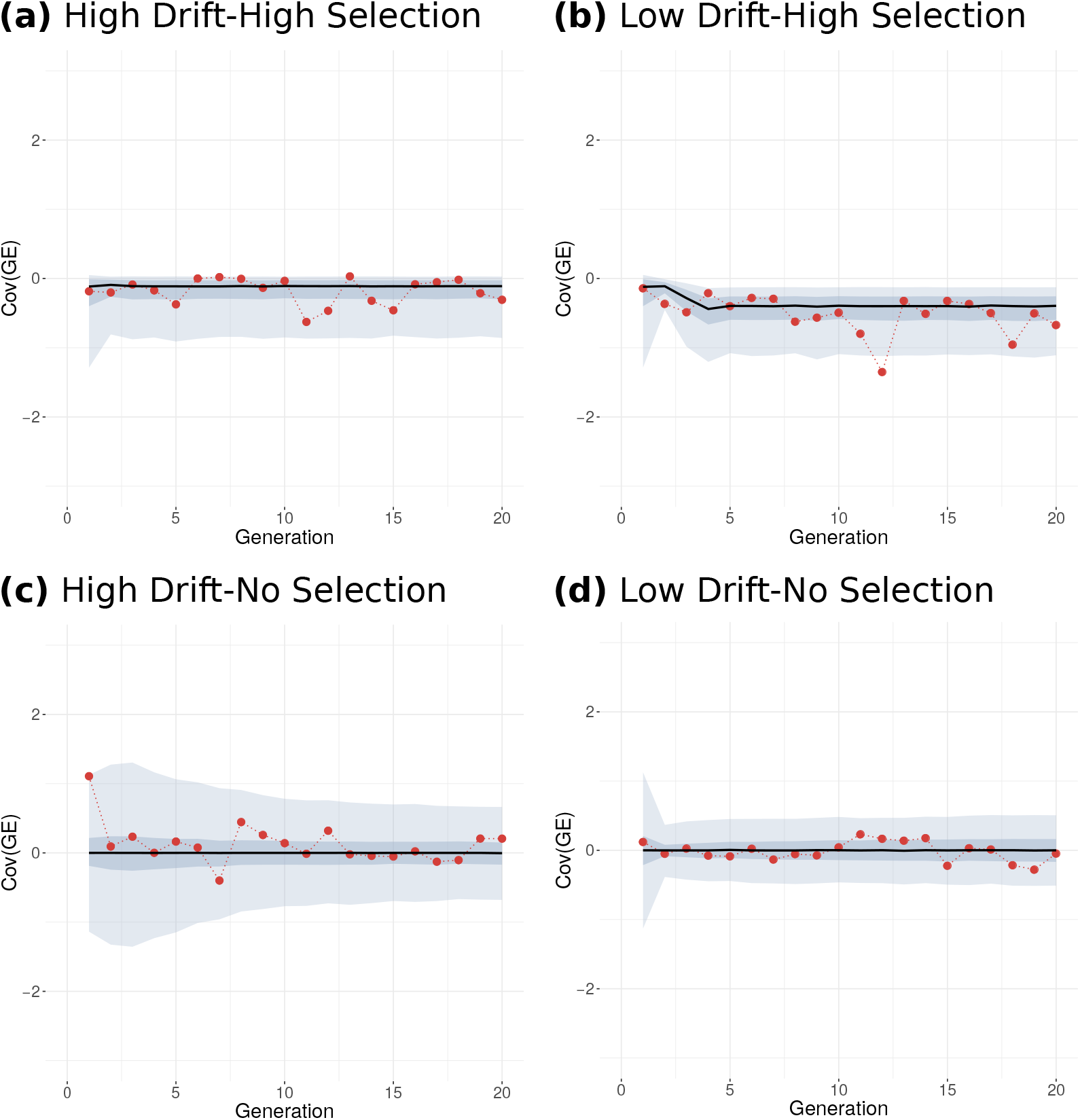
Evolution through time of the per-family covariance between environmental and genotypic values of the selected individuals, under our four simulated regimes. The black line represents the evolution of the median value over 2000 simulations in HDHS (a), LDHS (b), HDNS (c), LDNS (d). The shaded area corresponds to the 5^*th*^-95^*th*^ percentile (light blue) and to the 25^*th*^-75^*th*^ percentile (dark blue). One randomly chosen simulated family is represented with red dotted line, to highlight the inter-generation stochasticity. No significant autocorrelation was found.

**Figure S10.**
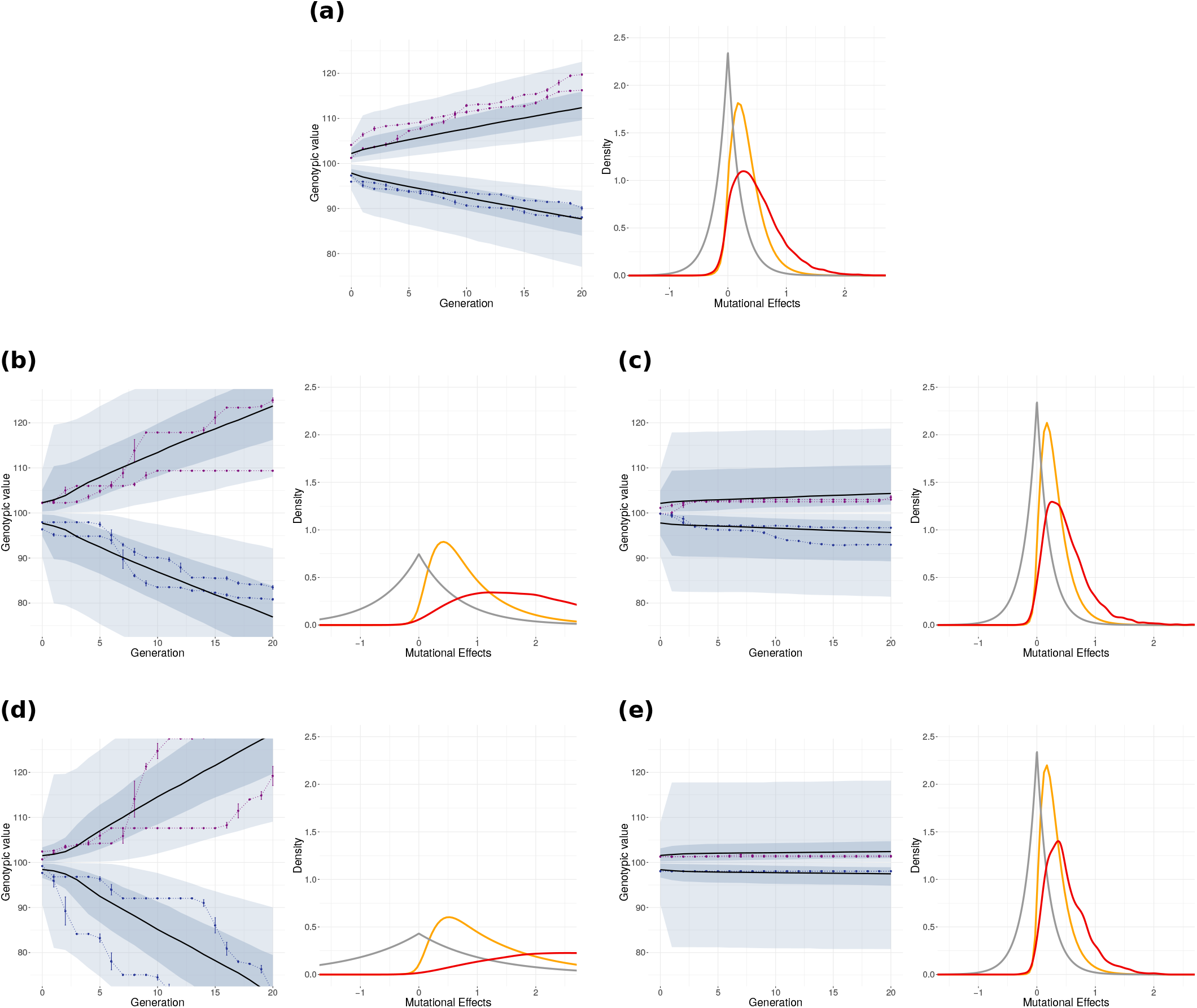
Comparison between simulated HDHS regime under various mutational parameters. Each panel (a) to (e) replicates Fig. 1 on the left and Fig. 4 on the right. The left side of each panel represents the mean genotypic values of the selected progenitors per family (expressed in Days To Flowering, DTF) across generations, violet/blue color identifies the late/ early population. In each population, the black line represents the evolution of the median value over 2000 simulations of the family genotypic mean. The shaded area corresponds to the 5^*th*^-95^*th*^ percentile (light blue) and to the 25^*th*^-75^*th*^ percentile (dark blue). In addition, two randomly chosen simulations are shown with dotted lines. The right side of each panel represents the distribution of effects of incoming *de novo* and fixed mutations under HDHS. Density distributions are shown for all incoming *de novo* mutational effects in grey — reflected exponential distribution —, and fixed mutations over 2000 simulations in red. Theoretical expectations from (Eq: 16) are plotted in gold. Panel (a) corresponds to the main text simulation parameters, so that HDHS: *U* = 1000 × 6000 × 30 × 10^−9^ = 0.18, *E*(λ_*M*_) = 4.69, 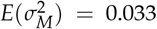. (b) and (c) correspond to simulations with 100 loci and the same mutation rate per locus *μ* = 6000 × 30 × 10^−9^ = 0.018, while (d) and (e) correspond to simulations with 1000 loci but a mutation rate per locus *μ* = 6000 × 1 × 10^−9^ = 0.006. Panels (b) and (d) were obtained with the same total mutational variance 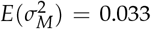 as HDHS but smaller *λ_M_*, (*E*(*λ_M_*) = 1.50 and *E*(*λ_M_*) = 0.86 respectively). (c) and (e) were obtained with the same mutational variance per locus as HDHS (*E*(*λ_M_*) = 4.69) but smaller total mutational variance than HDHS 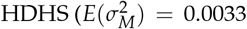 and 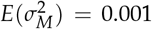 respectively). Overall we have :

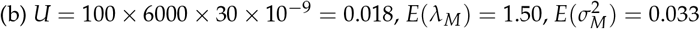

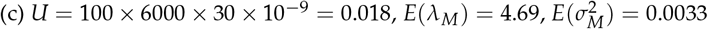

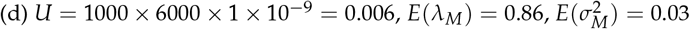

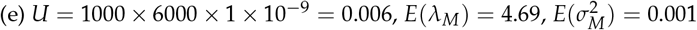

Note that while the distributions of incoming *de novo* and fixed mutational effects are similar among panels (a), (c), (e), the number of fixed mutations in (c) and (e) is much lower than in (a) accounting for the lack of selection response observed in those panels.

**Figure S11.**
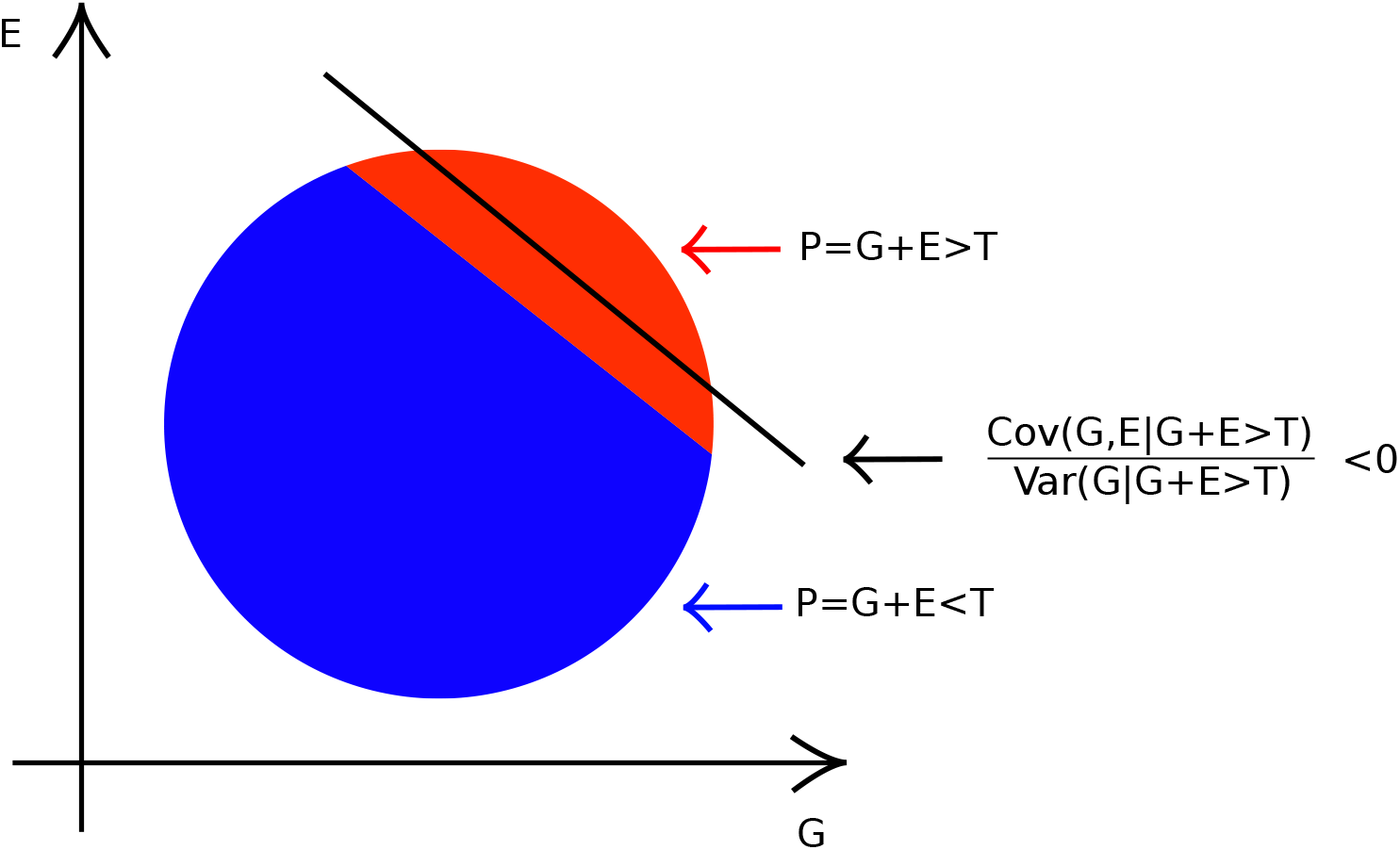
Schematic representation of the impact of selection on Cov(*G*, *E*). For illustration purposes, let P the sum of two independent and identically distributed random variables, G and E, such that both G and E follow a standard normal distribution, *i.e. P* = *G* + *E* with 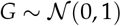 and 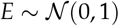. The black line represent the regression of *E*_|*selected*_ on *G*_|selected_ with a negative slope 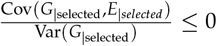.

**Figure S12.**
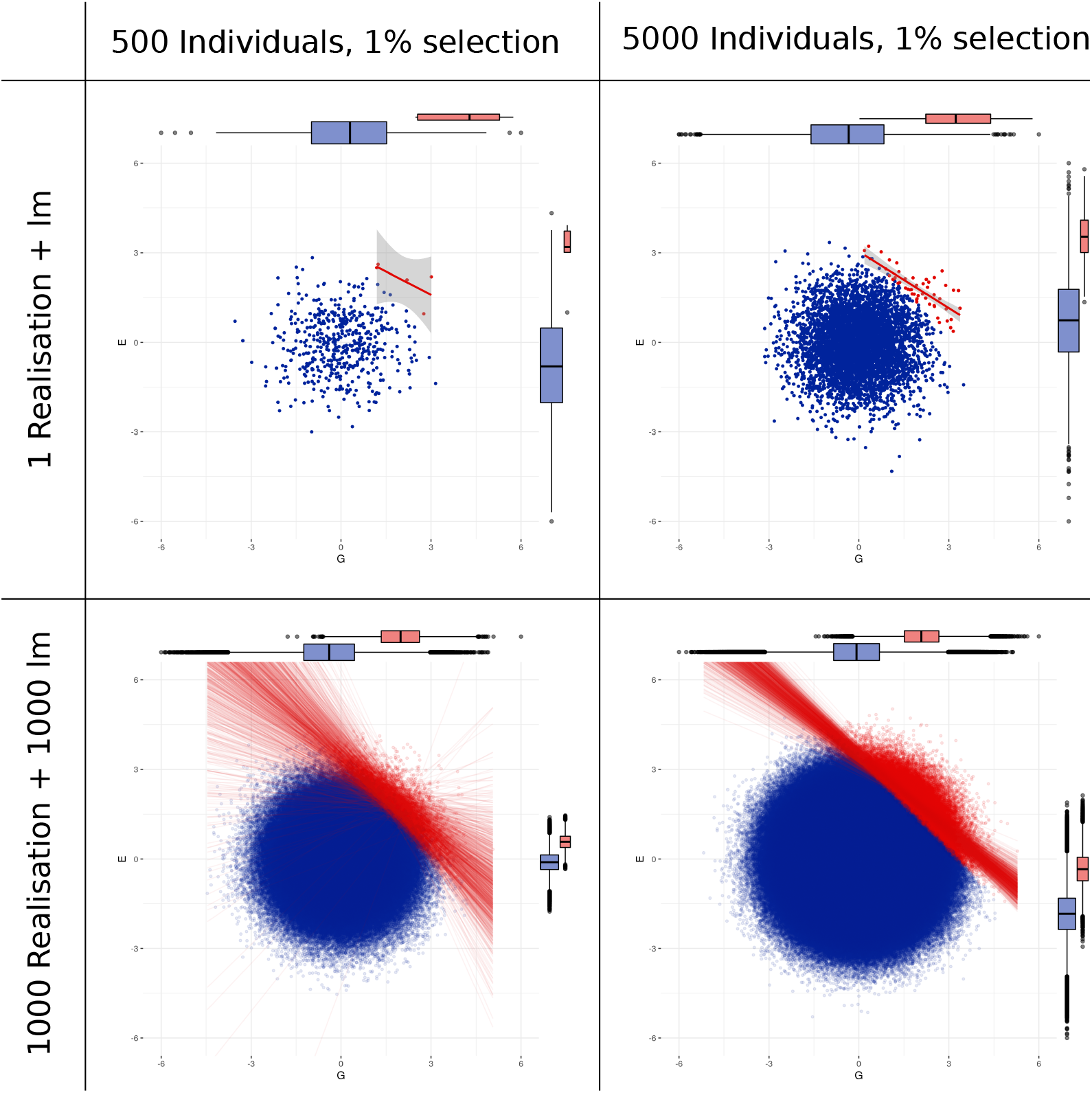
Schematic representation of the impact of selection and drift on Cov(*G*, *E*). Let P the sum of two independent random variables, G and E, such that both G and E follow a standard normal distribution, *i.e. P* = *G* + *E* with 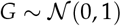 and 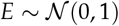. Let sample 500 individuals from P and plot *E* = *f* (*G*) (right columns), respectively 5000 (left columns) and select (red dots) the best 1% based on P. The upper row represents one realisation, with the red line corresponding to the regression of *E*_|selected_ on *G*_|selected_ with a negative slope 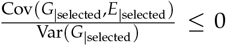. The lower row represents the realisation of 1000 independent sampling of 500 and 5000 individuals, with the corresponding linear regressions. We observe a lower lesser exploration of possible values (red plus blue area) under low population size and a high stochasticity in the values of Cov(*G*_|selected_, *E*_|selected_)

